# Plant kleptomaniacs: geographic genetic patterns in the amphi-apomictic *Rubus* ser. *Glandulosi* (Rosaceae) reveal complex reticulate evolution of Eurasian brambles

**DOI:** 10.1101/2024.01.16.575855

**Authors:** Michal Sochor, Petra Šarhanová, Martin Duchoslav, Michaela Konečná, Michal Hroneš, Bohumil Trávníček

## Abstract

*Rubus* ser. *Glandulosi* represents a unique model of geographic parthenogenesis on a homoploid (4x) level. We employed double digest restriction-site associated DNA sequencing (ddRADseq) in 587 individuals of different *Rubus* taxa and modelling of suitable climate to characterize evolutionary and phylogeographic patterns and shed light on the geographic differentiation of apomicts and sexuals. Six ancestral species were identified that contributed to the contemporary gene pool of *R.* ser. *Glandulosi*. While sexuals were introgressed from *R. dolichocarpus* and *R. moschus* in West Asia and from *R. ulmifolius* agg., *R. canescens* and *R. incanescens* in Europe, apomicts were characterized by alleles of *R.* subsect. *Rubus*. Gene flow between sexuals and apomicts was also detected, as well as occasional hybridization with other taxa. We hypothesize that sexuals survived the last glacial period in several large southern refugia, whereas apomicts were mostly restricted to southern France from whence they quickly recolonized Central and Western Europe. The secondary contact of sexuals and apomicts was probably the principal factor that established geographic parthenogenesis in *R.* ser. *Glandulosi*. Sexual populations are not impoverished in genetic diversity along their borderline with apomicts and maladaptive population genetic processes likely did not shape the geographic patterns.

**Highlights:**

- Geographic parthenogenesis in *Rubus* ser. *Glandulosi* is caused by secondary contact
- Reticulate evolution in Eurasian brambles is more extensive than expected
- At least six species contributed to the evolution of *Rubus* ser. *Glandulosi*
- Extinct ancestor’s gene pool is associated with apomixis

Graphical abstract

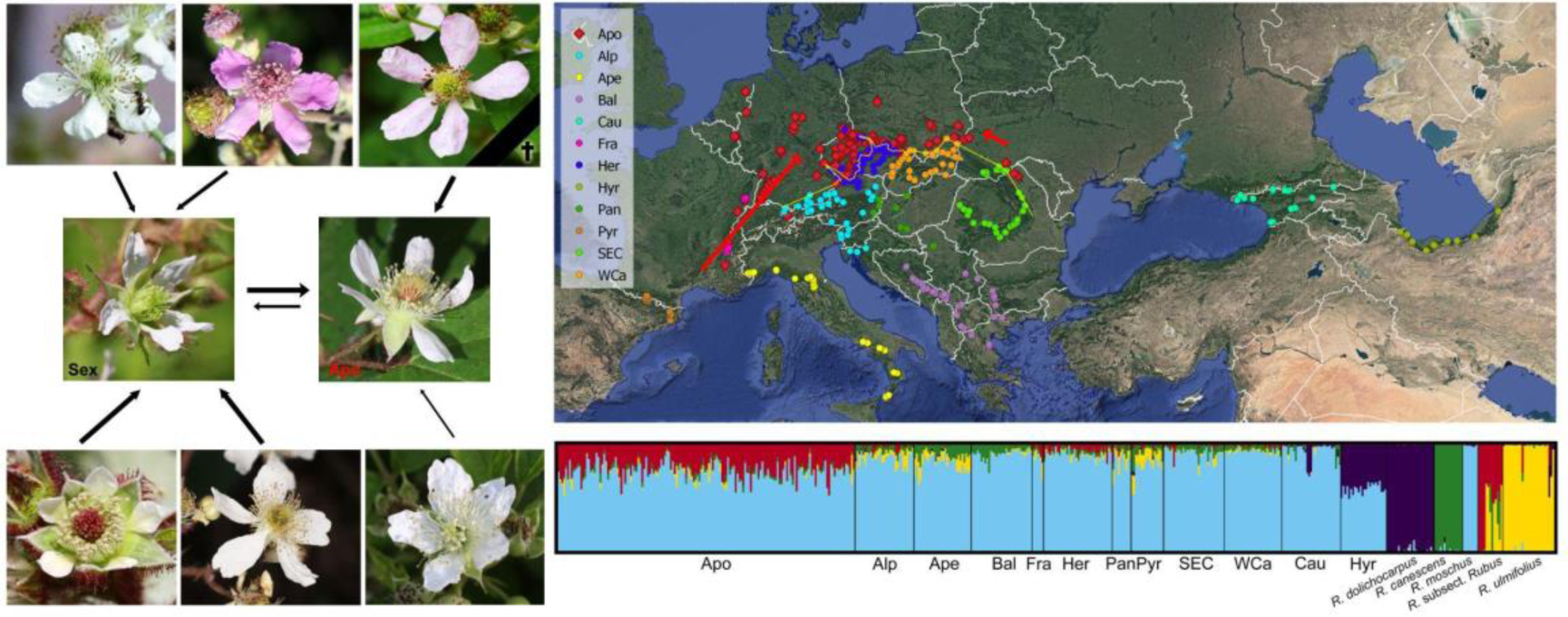

## 1. Introduction

*Rubus* subgen. *Rubus* (blackberries, brambles) is notorious for its complex evolutionary patterns, the main evolutionary driving forces being hybridization, polyploidization, and quaternary range contractions and expansions (Sochor et al., 2015, 2017). Most of the species are polyploid facultative apomicts, i.e., they are able to form seeds both sexually and asexually via gametophytic apomixis (Kurtto et al., 2010; Šarhanová et al., 2012, and references therein). Only five diploid species are extant in Eurasia, namely *R. ulmifolius* agg. (including *R. ulmifolius* Schott and *R. sanctus* Schreb.), *R. canescens* DC., *R. moschus* Juz., *R. dolichocarpus* Juz. and *R. incanescens* Bertol. (Gustafsson, 1943; Krahulcová et al., 2013; Kasalheh et al., under review). The first two species have a very large distribution from the Atlantic Ocean to Afghanistan (*R. ulmifolius* agg.) or to Armenia (*R. canescens*; Fig. 1). The former also shows strong inter-regional genetic differentiation, which is traditionally treated taxonomically on the species level: *R. ulmifolius* in the west and *R. sanctus* in the east (Monasterio-Huelin and Weber, 1996). The ranges of the other species are much smaller; *R. moschus* is endemic to the western Lesser Caucasus (Juzepczuk, 1925), *R. dolichocarpus* extends from the central Caucasus to eastern Hyrcania (Kasalkheh et al., in prep.), and *R. incanescens* is scattered along the French/Italian Mediterranean coast and some neighbouring areas (Kurtto et al., 2010). Furthermore, two tetraploid sexuals are known from the continent, namely a part of *R.* ser. *Glandulosi* (Wimm. et Grab.) Focke, and *R. caesius* L., both of which have a very large distribution (Figs 1, 2).

**Fig. 1:**
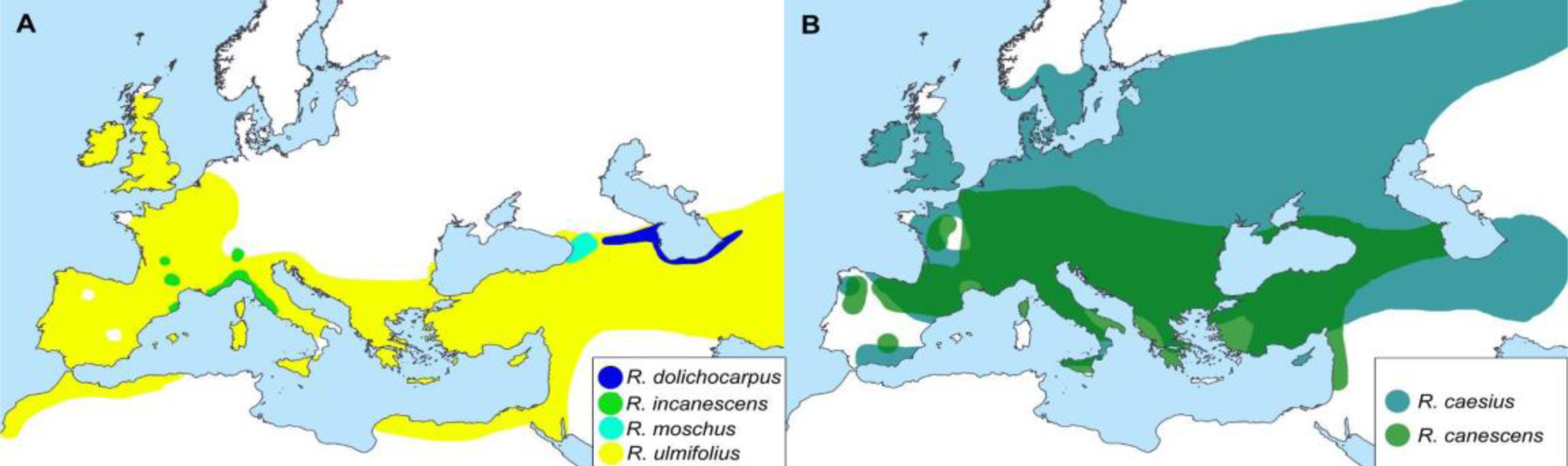
Approximate distributions of *Rubus* ancestors based on Kurtto et al. (2010), Juzepczuk (1941), and our sampling (Supplementary data 1). *Rubus idaeus* is not included.

**Fig. 2:**
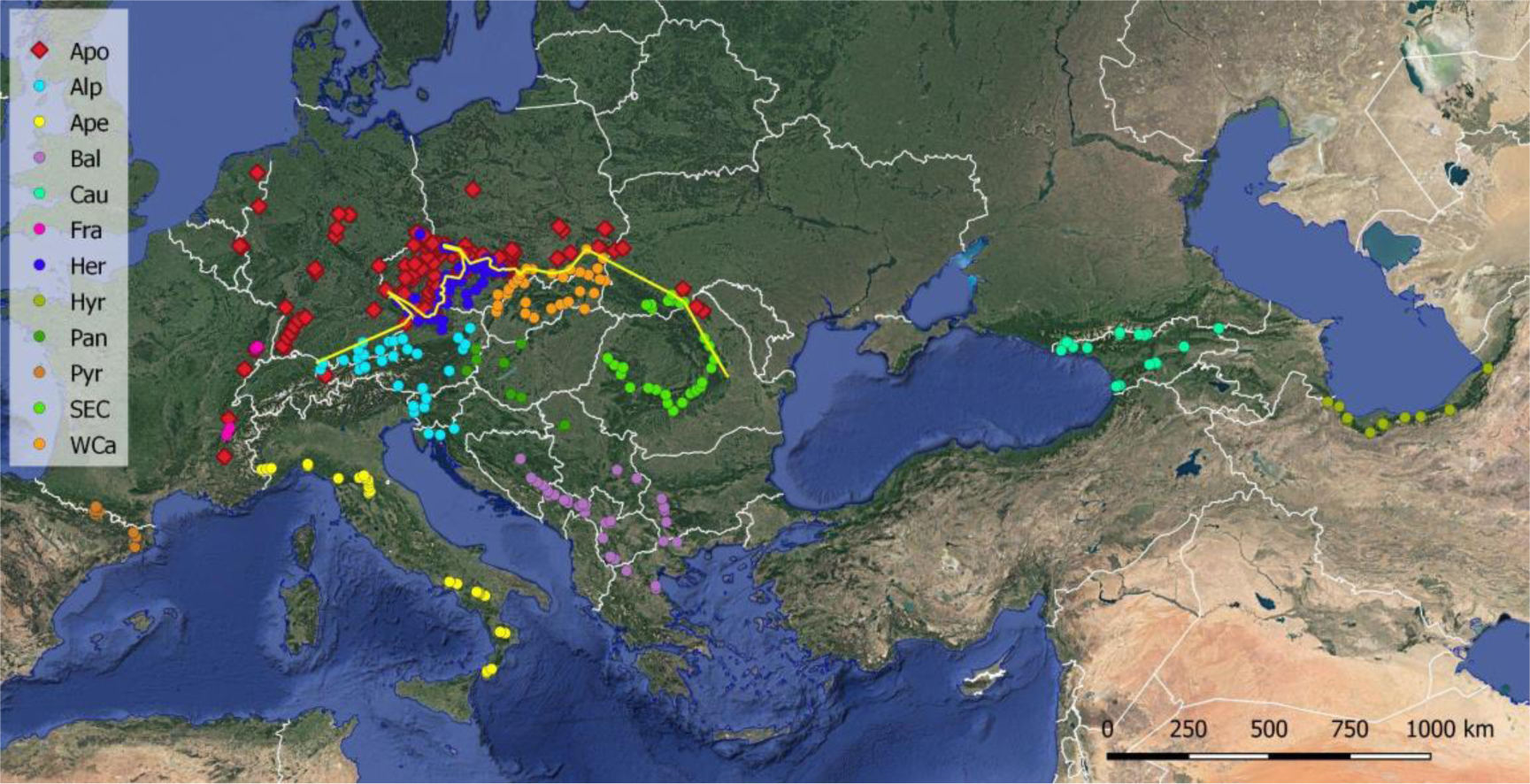
Map of sampled individuals of *Rubus* ser. *Glandulosi* with their population assignment: apomicts (Apo); sexuals are distinguished according to their geographic origin – Alps (Alp), Apennines (Ape), Balkans (Bal), France (Fra), Hercinia (Her), Pannonia (Pan), Pyrenees (Pyr), South-eastern Carpathians (SEC), Western Carpathians (WCa), Caucasus (Cau), Hyrcania (Hyr). The borderline between sexuals and apomicts is shown as a yellow line. Background layer: Google Earth.

All extant sexuals are known to have contributed in some way to the evolution of polyploid apomicts, with the exception of *R. incanescens*, whose alleles have not yet been detected in any apomict, although it can form triploid hybrids (Sochor et al., 2015). Furthermore, at least one extinct European and one Caucasian sexual have been inferred from phylogenetic studies. Alleles of the extinct European ancestor have been detected particularly in *R.* ser. *Nessenses* (whose other parent is *R. idaeus* L.) and *R.* ser. *Rubus* (i.e., *R.* subsect. *Rubus*, a group formerly called “*Suberecti*”), but to a lesser extent also in other series of high-arching brambles. This species was closely related to the erect North American taxa, such as *R.* ser. *Canadenses* (L.H. Bailey) H.E. Weber and *R.* ser. *Alleghenienses* (L.H. Bailey) H.E. Weber. (Sochor et al., 2015). The second extinct or unknown species with an unsuspected phenotype is presumed to be ancestral for the Caucasian polyploids (Sochor and Trávníček, 2016).

Forming the opposite extreme of the morphological continuum to *R.* subsect. *Rubus*, tetraploids of *R.* ser. *Glandulosi* are supposed to be the dominant ancestor of prostrate or low-arching brambles characterized by stipitate glands on stems (Sochor et al., 2015). They likely originated from *R. moschus* or its ancestor during the last or preceding interglacial periods. This taxonomically hitherto unresolved group exhibits a peculiar pattern of geographic parthenogenesis; whereas plants from North-western Europe and extra-Carpathian regions of Poland, Ukraine and Romania are facultative apomicts, populations from Southern and Central Europe, the Caucasus and Hyrcania are strictly sexual (Fig. 2). The distribution of genotypic diversity in apomicts implies a secondary contact between the reproductive groups. Consequently, phylogeography and neutral microevolutionary processes appear as the main factors propelling geographic parthenogenesis in this taxon (Sochor et al., under review). However, population genetic processes typical for small fragmented populations (Keller and Waller, 2002; Willi et al., 2013) may also play some role in geographic differentiation between reproductive modes. This concept presumes that marginal populations of sexuals are impoverished in genetic diversity, resulting in reduced adaptability and an inability to compete with apomicts, which are more resistant against genetic drift and inbreeding and outbreeding depression thanks to their fixed heterozygosity (elevated by hybridity and polyploidy) and asexuality (Haag and Ebert, 2003; Tilquin and Kokko, 2016; Sochor et al., 2017). In theory, this hypothesis applies for both primary and secondary contact zones.

Due to the unique pattern of geographic parthenogenesis and more-or-less apparent morphological deviations of some apomictic *Glandulosi* genotypes, we originally suspected that the apomicts are recently derived from sexual populations either directly, or via hybridization with some other (i.e., non-*Glandulosi*; see Table 1), apomictic taxon. Such apomictic taxa, which could potentially pose a source of apomixis for the newly formed *Glandulosi* apomicts, occur in the entire range of sexuals, including the Caucasus and Hyrcania (Kurtto et al., 2010; Sochor and Trávníček, 2016; Kasalkheh et al., in prep). Alternatively, the *Glandulosi* apomicts may have been formed already before the last glaciation, i.e., before the establishment of current geographic and genetic patterns in the European *R. subgen. Rubus*. These two scenarios—Pleistocene vs. Holocene origin of the *Glandulosi* apomicts—are expected to result in different patterns of allele sharing between the *Rubus* ancestral taxa and the *Glandulosi* apomicts.

**Table 1:**
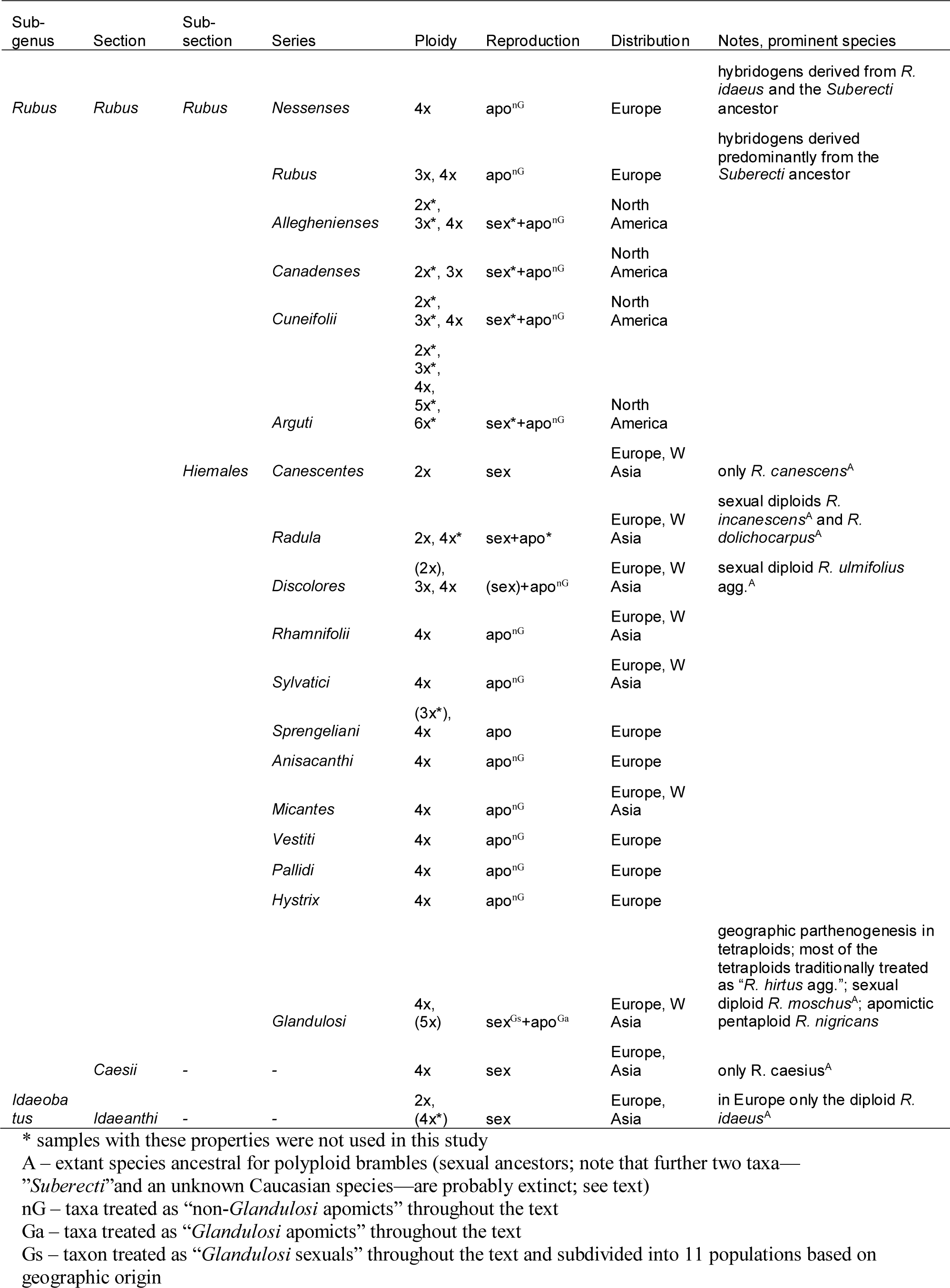
Taxonomic overview over taxa used in this study with ploidy levels, reproductive mode, distribution and other notes compiled from Thompson (1997), Kurtto et al. (2010), Krahulcová et al. (2013), Sochor et al. (2015), and van de Beek (2021).

However, the phylogenetic and phylogeographic patterns or structuring of genetic diversity have not yet been characterized due to the lack of polymorphism in the commonly used phylogenetic markers (Sochor et al., 2015) and the absence of high-resolution genomic data in *R.* ser. *Glandulosi*. Taking into consideration all the peculiarities of *R.* ser. *Glandulosi*, we aim to resolve genetic geographic patterns in the group across its range. Specifically, we aim to (1) test the role of geographic structure of sexuals’ genetic diversity in the formation of geographic parthenogenesis patterns, (2) reconstruct the roles of other *Rubus* taxa in the evolution of *R.* ser. *Glandulosi*, (3) explain the origin of the *Glandulosi* apomicts, and (4) identify the glacial refugia and postglacial recolonization routes of both sexuals and apomicts of *R.* ser. *Glandulosi*. Ultimately, we evaluate the importance of phylogeography in the formation of geographic parthenogenesis.

## 2. Materials and Methods

### 2.1. Sampling

Due to the patterns of geographic parthenogenesis, our sampling was focused on tetraploid members of *R.* ser. *Glandulosi* across its distribution range. For this taxon, we sampled individuals analysed in our preceding work (Sochor et al., under review). As complex evolutionary patterns were expected between this taxon and other *Rubus* species, our sampling additionally included other taxa of *R.* sect. *Rubus* and their two known ancestors outside this section—*R. caesius* and *R. idaeus* (Table 1). Due to the instability of nomenclature and some questionable recent nomenclatural changes, we follow Kurtto et al. (2010) for taxa names. *Rubus* ser. *Glandulosi* (or *Glandulosi* hereafter) samples were divided into 12 groups (hereafter termed “populations”) according to their reproductive mode as assessed by the flow-cytometric seeds screen and/or SSR genotyping (Sochor et al., under review) and their geographic origin (Fig. 2). As all apomictic individuals available to us have been genotyped by SSR (Sochor et al., under review), usually one individual per genotype was selected for sequencing. In total, 159 individuals (152 genotypes) were included in the single apomictic population (apo), covering tetraploids, additionally the common pentaploid *R. nigricans* Danthoine (syn. *R. pedemontanus* Pinkw.; one genotype with two individuals) and three other pentaploid genotypes from south-eastern France. Tetraploid sexual individuals were divided into 11 populations: the Alps (Alp, 31 individuals), Apennines (Ape, 30 ind.), Balkans (Bal, 32 ind.), the French mountains of Chartreuse and Vosges (Fra, 6 ind.), Hercynia (Her, 36 ind.), Pannonia (Pan, 10 ind.), Pyrenees (Pyr, 17 ind.), Southern and Eastern Carpathians (SEC, 32 ind.), Western Carpathians (WCa, 30 ind.), Caucasus (Cau, 31 ind.), and Hyrcania (Hyr, 24 ind.). Besides the twelve groups from *R.* ser. *Glandulosi*, all extant Eurasian diploid species were included in the study, sampled across their entire ranges wherever possible: *R. ulmifolius* agg. (27 ind.), *R. canescens* (15 ind.), *R. moschus* (8 ind.), *R. dolichocarpus* (25 ind.), *R. incanescens* (4 ind.), *R. caesius* (15 ind.), *R. idaeus* (5 ind.), and an outgroup of 50 individuals of non-*Glandulosi* apomictic taxa (mostly recognized microspecies) selected to cover most of the other series of *R.* sect. *Rubus* (Table 1). For the complete list of analysed specimens, see Supplementary data 1.

### 2.2. Whole genome sequencing and assembly

Due to the lack of whole-genome reference sequences for the studied group, a *R. moschus* specimen (MS36/14Bs from Adjara, Georgia) was shot-gun sequenced using two approaches. First, long reads were generated by Oxford Nanopore Technologies (ONT) using Rapid Sequencing kit SQK-RAD004 and the MinION sequencing device and two R9.4.1 flow cells following the manufacturer’s instructions. The genomic DNA was extracted using Invisorb Spin Plant Mini Kit (Invitec Molecular, Berlin) and subsequently size-selected for fragments of >40 kbp by Short Read Eliminator XL Kit (Circulomics). Basecalling was performed in the software MINKNOW 21.02.1 using the DNA High-Accuracy algorithm. Second, short high-accuracy reads were generated from the same specimen by Macrogen Europe (Amsterdam) on the Illumina Novaseq6000 sequencing platform in the 2×150 bp configuration, using the TruSeq DNA PCR-Free kit with a 350 bp insert for library preparation.

The ONT data were checked in FastQC 0.11.9 (Andrews, 2010) and adaptor sequences were trimmed in LongQC 1.2.0c (Fukasawa et al., 2020). NANOFILT 2.8.0 (De Coster et al., 2018) was used for filtering data on the minimum read length of 150 bp, minimum average read quality score of 5, and 10 nucleotides were trimmed from start of each read. Whole-plastome sequence was assembled from the Illumina data in GETORGANELLE 1.7.6.1 (Jin et al., 2020) using kmer sizes of 21, 45, 65, 85 and 105 and the embryophyta plant plastome database.

Completeness of the sequence was checked visually in alignment with publicly available *Rubus* plastomes. ONT sequences that mapped on the plastome sequence were subsequently filtered out via mapping in MINIMAP2 (ver. 2.24; Li, 2018) and only non-plastome reads were used for *de novo* genome assembly in SMARTDENOVO (Liu et al., 2021) with default parameters and kmer size set to 19. Subsequently, the sequence was polished in MEDAKA 1.6.0. (https://github.com/nanoporetech/medaka) using the base-called ONT reads, and in NEXTPOLISH 1.3.0 (Hu et al., 2019) using the Illumina data. Finally, the contigs were scaffolded into chromosomes in RAGTAG 2.1.0 (Alonge et al., 2022) using its scaffold function and two reference genome sequences: *R. ulmifolius* ‘Burbank Thornless’ genome ver. 1 (https://www.dnazoo.org/assemblies/Rubus_ulmifolius) and *R. occidentalis* genome ver. 3 (VanBuren et al., 2018). BUSCO analysis (ver. 5.4.3; Manni et al., 2021) was performed with 2,326 BUSCO groups of the Eudicots database. All the computationally demanding calculations were performed on the Czech National Grid Infrastructure MetaCentrum.

### 2.3. Double digest restriction-site associated DNA sequencing (ddRADseq.)

Genomic DNA was extracted from silica gel-dried leaves by CTAB method (Doyle and Doyle, 1987) and purified by Mag-Bind® TotalPure NGS (Omega Bio-tec), or using Invisorb Spin Plant Mini Kit (Invitec Molecular, Berlin). In both cases, it was finally purified by precipitation with sodium acetate (concentration of 0.3 M in the DNA isolate) and ethanol (100%, double of the DNA volume). The quality of DNA was checked using 1.5% agarose gel electrophoresis and concentration was measured by Qubit 4 fluorometer with 1X dsDNA HS Assay Kit (Thermo Fisher Scientific). As the basis for ddRAD library protocol development, the protocol of Peterson et al. (2012) was utilized. First, 100 ng of DNA was cleaved with PstI HF and MseI restrictases in CutSmart® buffer (New England Biolabs; 3 hours at 37°C). P1 and P2 barcoded adapters corresponding to the restriction site of the enzymes were ligated immediately afterwards with T4 ligase (NEB) at 16°C overnight. The enzymes were heat-killed with 65°C for 10 minutes. All samples with the same P2 and different P1 adapters were pooled, purified by Mag-Bind kit, and size-selected with Pippin Prep 1.5% agarose gel cassette (Sage Science) for fragment sizes of 250–500 bp. PCR-enrichment was performed in 18 cycles in ten 10μL reactions per sublibrary with Phusion® HF PCR Mastermix (New England BioLabs) by mixing 5µl 2× Phusion Master mix, 0.4 µM of each standard Illumina P5 (5′-AATGATACGGCGACCACCGAGATCT-3′) and P7 (5′-CAAGCAGAAGACGGCATACGAGAT-3′) PCR primers, 3.2 µl H2O, and 1µl restricted-ligated DNA pool. Each sublibrary was purified with Mag-Bind kit or 1.2× SPRIselect beads (Beckman Coulter) and quantified on Qubit. All sublibraries were pooled equimolarly and sequenced on NovaSeq 6000 platform (Illumina) using the SP reagent kit v1.5 in 2×150 bp configuration at the Institute of Experimental Botany, Czech Academy of Sciences, Olomouc.

### 2.4. ddRADseq data analysis

Raw reads were checked for quality in FASTQC 0.11.9 (https://www.bioinformatics.babraham.ac.uk/projects/fastqc) and demultiplexed in STACKS 2.63 (Catchen et al., 2013). SEQPURGE 2019_09 (Sturm et al., 2016) was used for quality filtering and adapter trimming with default settings. Reads were subsequently mapped to the reference whole-genome sequence of *R. moschus* using BWA-MEM algorithm (bio-bwa.sourceforge.net) and converted to BAM files via SAMTOOLS (Li et al., 2009). A catalogue of loci was created in the GSTACKS module of STACKS and was subsequently used for data filtering in the POPULATIONS module. Alternative *de novo* and reference-based pipelines with tetraploid settings were also tested in IPYRAD (Eaton and Overcast, 2020), but these resulted in less recovered loci and/or more noisy phylogenetic signal (not shown), and were, therefore, not used further.

For *R.* ser. *Glandulosi* populations and ancestral taxa, diversity indices (heterozygosity, F_IS_, number of private alleles) were calculated in POPULATIONS with the R parameter (minimum proportion of individuals to process a locus) set to 0.5. To assess correlation between genetic diversity indices and the distance from the borderline between sexuals and apomicts (sex-apo borderline sensu Sochor et al., under review; Fig. 2), the distance was measured for each sexual individual using the Distance-to-nearest hub function in QGIS 3.30.0 (www.qgis.org) and expected heterozygosity (H_exp_) was calculated for each individual sample in POPULATIONS (R=0.7, min-maf=0.05). Linear regression was subsequently calculated in NCSS 9 (www.ncss.com). As H_exp_ was significantly correlated with data coverage on the individual level, linear regression was also calculated with residuals of H_exp_ from its regression on the number of the recovered variant sites. Populations Fra, Pyr, Ape, Cau and Hyr were excluded from these analyses due to unclear placement of the borderline in Western Europe and geographical isolation of the Asian populations.

Private alleles were identified in POPULATIONS for each of the potentially ancestral species (diploids and *R. caesius*; Table 1) and subsequently traced using our custom-made script (available at github.com/hajnej/population_private_alleles) in the target (derived) individuals in populations.sumstats.tsv generated by another POPULATIONS run. Numbers of private alleles detected in each individual were divided by the individual’s total number of variant sites to avoid the effects of unequal coverage and missing data. Alleles identified as private for *R.* subsect. *Rubus* against the extant potentially ancestral species were considered as originating in the extinct *Suberecti* ancestor; these alleles were traced separately due to the hybrid origin of the *R.* subsect. *Rubus* polyploids.

To obtain another proxy for allele sharing between ancestral and derived taxa, a reference sequence for each potentially ancestral taxon (Table 1) was constructed from ddRADseq reads (hereafter termed “pseudoreference” to avoid confusion with *R. moschus* whole-genome reference sequence); reads from the ancestral taxa were used for *de novo* assembly in IPYRAD (clustering threshold 0.9, other parameters left default), consensus sequences of all samples within species were mixed together and microbial contaminants were excluded by BBDUK (sourceforge.net/projects/bbmap) by mapping to the available viral, bacterial, fungal and protozoan reference genomes from the NCBI Reference Sequence Database. Reads of each individual of the derived taxa were mapped by BWA-ALN (bio-bwa.sourceforge.net) with *maxDiff* set to one, and proportion of mapped reads was calculated in GSTACKS as a proportion of all reads that were not unmapped (i.e., including those with low mapping qualities and soft-clipped alignments). To correct for inter-individual variation in data quality (and thus read mapping), the average mapping efficiency across pseudoreferences within an individual was subtracted from each value.

STRUCTURE 2.3.4 (Pritchard et al., 2000) was used for inferring population genetic structure and ancestry for two sample sets: (1.) *Basic set* was composed of all samples except *R. incanescens* (due to small sample size), *R. idaeus*, *R. caesius* and its three hybrid derivatives (due to their apparently negligible role in the evolution of *R.* ser. *Glandulosi*, and to reduce complexity of the data set), and polyploid outgroups (for their complex origin), but included *R.* ser. *Rubus* and North American taxa (representatives of *R.* subsect. *Rubus*); and (2.) the *European set* further excluded all extra-European samples. The following setup was used for the computation: admixture model, no linkage, no prior population information, K in the range 3–7 in ten replicate runs for each K with 150,000 burn-in iterations followed by further 200,000 MCMC iterations. STACKS parameters for the input data filtering were set as follows: R=0.7, min-maf=0.05, one random SNP per locus. The best K was selected using the method of Evanno et al. (2005) as implemented in STRUCTURE HARVESTER (Earl and vonHoldt, 2012), as well as according to similarities among runs and interpretability of the results. CLUMPAK (Kopelman et al., 2015) was used for the results post-processing and visualization.

RADPAINTER and FINERADSTRUCTURE 0.3.3 (Malinski et al., 2018) analyses were performed with another two sample sets: (3.) the *Complete sample set* (including all polyploid and diploid outgroups) and (4.) the *European Glandulosi set* (i.e., excluding all outgroups and Asian *Glandulosi* samples, as well as 25 samples with >80% of missing data). The analyses included calculation of the coancestry matrix, assignment of individuals into clusters (admixture model with 50,000 or 250,000 burn-in for the two datasets, respectively, followed by 100.000 or 500.000 MCMC iterations with the thin interval of 1000) and tree building (with the same number of iterations). The associated plotting R script FINERADSTRUCTUREPLOT.R was used for generating coancestry heatmaps. POPULATIONS parameters for the input data filtering were set at R=0.5, min-maf=0.05.

### 2.5. Present-day and paleoclimatic distribution modelling

Models of suitable climate were computed for the tetraploid *R.* ser. *Glandulosi* using present-day bioclimatic variables from WorldClim 1.4 (Hijmans et al., 2005), with a spatial resolution of 2.5 arc minutes. Due to the large overlap of ecological niches of tetraploid sexuals and apomicts (Sochor et al., under review), the two reproductive groups were treated together as a single taxon. As the occurrence records were spatially biased towards Central Europe, occurrences closer than a minimum distance of 30 km to each other were removed, using SPTHIN library (Aiello-Lammens et al., 2015), and the resulting 192 occurrences were used in the modelling.

Because strong collinearity between bioclimatic variables may reduce model predictive ability (Svenning et al., 2011), confuse model interpretation (Baldwin, 2009), and reduce model transferability (Warren et al., 2014), we used a combination of selecting bioclimatic variables relevant to the ecological and physiological processes determining species distribution with a reduction of the number of variables through a statistical analysis. To assess multicollinearity between variables, 10,000 random points were generated within the minimal convex polygon around the thinned records plus a buffer of 2 arc minutes to extract cell values for all the variables. The resulting matrix was analysed using the vifcorr function from the USDM library (Naimi et al., 2014) to find strongly correlated pairs of variables. If the Pearson correlation coefficient was higher than |0.85| for a pair of variables, only one of them with a supposed tighter relationship with plant ecology was retained. At the end, six bioclimatic variables were selected: Bio2 (Mean diurnal temperature range), Bio3 (Isothermality), Bio5 (Maximum Temperature of Warmest Month), Bio6 (Minimum Temperature of Coldest Month), Bio12 (Annual Precipitation), and Bio15 (Precipitation seasonality).

Occurrence records and environmental data were used to calibrate the distribution model using the MAXENT method (Phillips and Dudík, 2008). This machine-learning method was selected because it generally performs well under different scenarios when only presence data are available (Elith et al., 2006). To build a suite of candidate models with differing constraints on complexity and to quantify their performance, MAXENT models were built and evaluated using the semi-automated streamline analysis in WALLACE 2.1 (Kass et al., 2023), which calls internally MAXNET 0.1.4 (cran.r-project.org/web/packages/maxnet/) and ENMEVAL 2.0 (Kass et al., 2021). Models used 10,000 background points generated within the buffer area of 2 arc minutes around thinned records to extract cell values for the selected variables. We applied a spatial data partitioning scheme (Checkerboard 2 with k = 4 groups; Muscarella et al., 2014), split occurrence data into training and validation subsets, tested all feature classes and their combinations, and a range of regularization multipliers (from 0.5 to 4.0 with 0.5 step value), and clamping procedure was accounted for (Phillips et al., 2009). To find the best model, we used primarily delta AICc (Warren and Seifert, 2011), but also estimated other evaluation metrics calculated by WALLACE 2.1: AUC for both training and validation subsets, AUCdiff (differences between the training and validation AUCs), and the omission rate (OR10pct, i.e. 10pct is the lowest suitability score for such localities after excluding the lowest 10% of them).

Final model computed for present-day climate was run in the standalone MAXENT application V.3.4.4 (Phillips and Dudík, 2008) and used a 10-fold cross-validation procedure and a clog-log output. The final model was projected to two paleoperiods: LGM (22,000 years BP) and mid-Holocene (6,000 years BP). The same bioclimatic variables and resolution as used for present-day climate were used for both paleoperiods, and variables were taken from two paleoclimatic models (CCSM4; Gent et al., 2011, and MIROC-ESM; Watanabe et al., 2011) of WorldClim 1.4 downscaled paleoclimate (Hijmans et al., 2005). Final maps were constructed in QGIS, using Google Satellite as a background map. Masks of LGM ice sheets and glaciers were extracted from Last Glacial Maximum version 1.0.1 compiled by Zentrum für Baltische und Skandinavische Archäologie (ZBSA; <u>zbsa.eu/en/last-glacial-maximum</u>), those of LGM continuous and discontinuous permafrost from Lehmkuhl et al. (2020) and LGM paleocoastlines from Zickel et al. (2016).

## 3. Results

### 3.1. Rubus moschus whole genome reference sequencing

Illumina sequencing provided 408.8 mil. reads (61.7 Gbp of total length, Q20 of 97.6%). After plastome and low-quality reads filtering, 1.54 mil. ONT reads (8.1 Gbp of total length, N50 = 9655 bp) were retained. The *de novo* assembly after polishing resulted in 2,170 contigs (plus one plastome contig) with a total length of 314,423,542 bp and N50 of 326,641 bp. The scaffolded assembly contained seven chromosome contigs (290.6 Mbp in total) and 511 unplaced contigs (24.0 Mbp), N50: 39,737,832 bp. BUSCO analysis identified 97.6% of complete (89.4% single-copy, 8.2% duplicated), 0.6% of fragmented, and 1.8% of missing genes. The final sequence is available on request from the corresponding author.

### 3.2. Genetic diversity and its spatial structuring

Among the *Glandulosi* populations, no marked differences were detected in both expected and observed heterozygosity (Table 2). F_IS_ index was influenced only by sample sizes. For example, the apomictic population exhibited markedly higher F_IS_, but after a random selection of 30 samples, it dropped to the value only slightly higher than those of the sexual populations of the same size (F_IS_=0.078) with only a negligible change in H_obs_ and H_exp_ (0.030 and 0.047, respectively). The absolute number of private alleles was also associated with sample size, but after correction by the number of samples per locus (PA/N per locus), the values varied only from 191 to 347 in European populations, while the Asian populations Cau and Hyr showed values of 809 and 795, respectively.

**Table 2:** Population genetic statistics based on 30,078 loci with 561,810 variant sites. N: number of individuals, H_obs_: observed heterozygosity, H_exp_: expected heterozygosity, F_IS_: inbreeding coefficient, SD: standard deviation of the preceding statistics, PA: number of private alleles.

| | N | N per locus | SD | $H_{obs}$ | SD | $H_{exp}$ | SD | $F_{IS}$ | SD | PA | PA/N per locus |
| --- | --- | --- | --- | --- | --- | --- | --- | --- | --- | --- | --- |
| Apo | 159 | 104.0 | 21.0 | 0.028 | 0.067 | 0.048 | 0.103 | 0.154 | 0.284 | 23,679 | 227.7 |
| Alp | 31 | 22.5 | 4.3 | 0.030 | 0.079 | 0.043 | 0.101 | 0.067 | 0.218 | 6,245 | 277.4 |
| Ape | 30 | 23.7 | 3.9 | 0.031 | 0.079 | 0.045 | 0.102 | 0.070 | 0.222 | 6,625 | 279.7 |
| Bal | 32 | 25.6 | 4.1 | 0.030 | 0.079 | 0.041 | 0.099 | 0.063 | 0.212 | 8,654 | 338.1 |
| Fra | 6 | 4.5 | 1.1 | 0.031 | 0.111 | 0.036 | 0.108 | 0.023 | 0.159 | 659 | 145.8 |
| Her | 36 | 23.4 | 4.7 | 0.028 | 0.077 | 0.041 | 0.100 | 0.066 | 0.215 | 4,379 | 187.0 |
| Pan | 10 | 6.8 | 1.8 | 0.029 | 0.092 | 0.041 | 0.107 | 0.041 | 0.193 | 2,200 | 325.2 |
| Pyr | 17 | 10.3 | 2.4 | 0.030 | 0.086 | 0.044 | 0.109 | 0.054 | 0.210 | 1,970 | 190.9 |
| SEC | 32 | 20.2 | 4.3 | 0.027 | 0.074 | 0.041 | 0.099 | 0.070 | 0.227 | 7,016 | 346.9 |
| WCa | 30 | 22.5 | 3.7 | 0.029 | 0.080 | 0.040 | 0.099 | 0.057 | 0.203 | 5,532 | 246.0 |
| Cau | 31 | 21.0 | 3.9 | 0.028 | 0.078 | 0.039 | 0.097 | 0.058 | 0.209 | 17,021 | 808.7 |
| Hyr | 24 | 15.7 | 3.7 | 0.028 | 0.082 | 0.041 | 0.107 | 0.057 | 0.208 | 12,498 | 795.4 |
| *R. caesius* | 15 | 7.2 | 3.3 | 0.045 | 0.130 | 0.066 | 0.149 | 0.066 | 0.241 | 29,650 | 4140.9 |
| *R. canescens* | 15 | 10.4 | 3.3 | 0.016 | 0.064 | 0.027 | 0.092 | 0.038 | 0.178 | 7,002 | 672.1 |
| *R. dolichocarpus* | 25 | 14.6 | 5.4 | 0.012 | 0.052 | 0.025 | 0.089 | 0.044 | 0.186 | 12,081 | 828.9 |
| *R. idaeus* | 5 | 3.0 | 1.3 | 0.016 | 0.091 | 0.023 | 0.095 | 0.024 | 0.155 | 21,451 | 7073.5 |
| *R. incanescens* | 4 | 3.7 | 0.7 | 0.019 | 0.108 | 0.017 | 0.081 | 0.003 | 0.107 | 5,678 | 1534.7 |
| *R. moschus* | 8 | 5.3 | 1.7 | 0.012 | 0.064 | 0.020 | 0.083 | 0.022 | 0.141 | 2,866 | 536.8 |
| *R. subsect. Rubus* | 14 | 9.8 | 3.2 | 0.039 | 0.102 | 0.063 | 0.132 | 0.079 | 0.255 | 36,822 | 3753.6 |
| *R. ulmifolius* agg. | 27 | 20.1 | 6.7 | 0.021 | 0.064 | 0.042 | 0.108 | 0.077 | 0.240 | 19,281 | 961.6 |

There was no correlation between the genetic diversity (H_exp_) of sexual individuals and their distance from the sex-apo borderline when analysed directly. However, a significant negative correlation was found when using residuals of H_exp_ from its regression on the number of variant sites (Fig. 3).

**Fig. 3:**
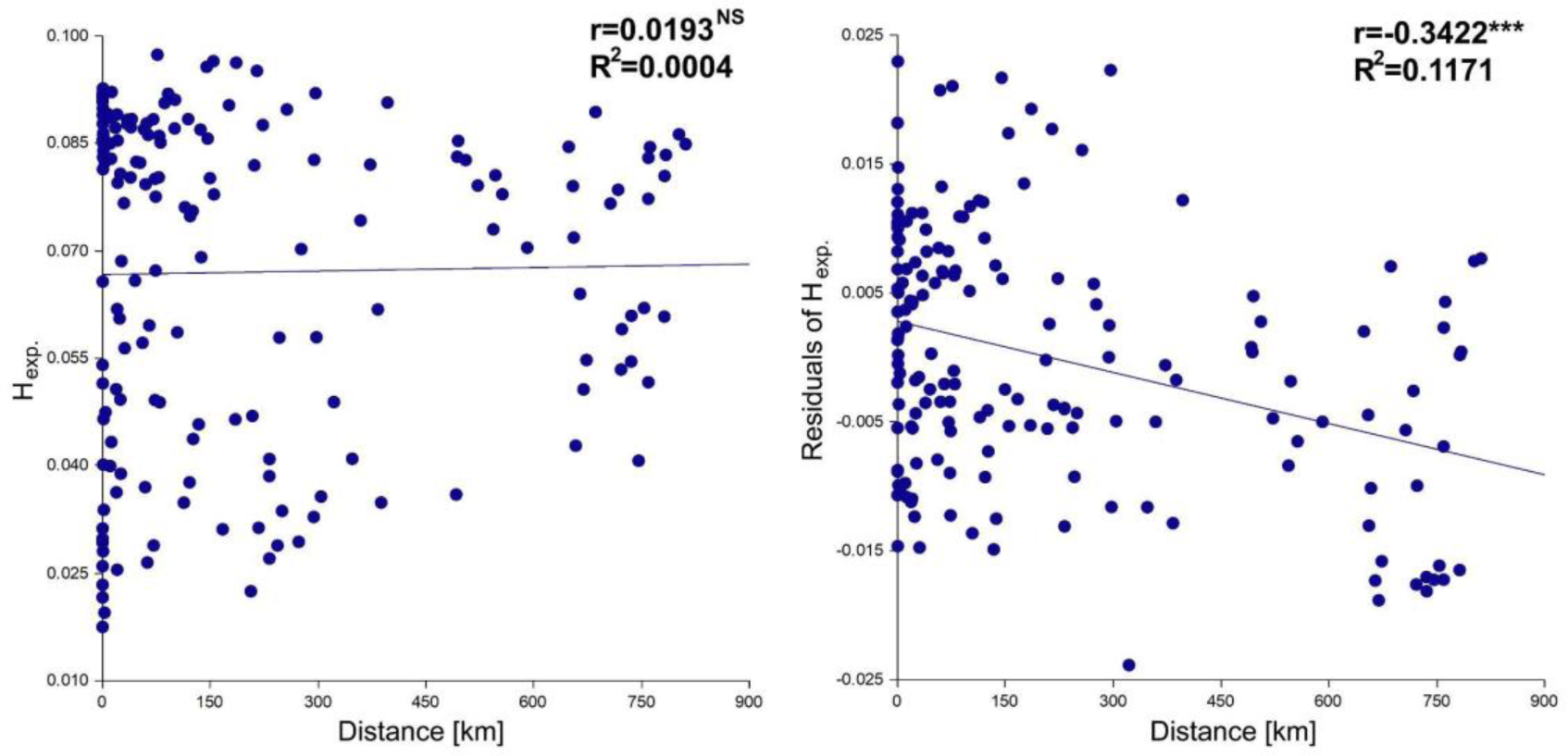
Linear regression of expected heterozygosity of individual genotypes (H_exp_; left) or its residuals (from linear regression of H_exp_ on the number of recovered variant sites; right) on the distance from the sex-apo borderline (Distance); only Central and Southeast European sexual individuals (excluding Fra, Pyr, Ape, Cau, and Hyr) with >6,000 variant sites were included (169 individuals); the analysis is based on 15,581 loci with 19,643 variant sites.

### 3.3. Private allele tracing and mapping of reads to pseudoreferences

Although the read mapping generally provided a somewhat weaker signal, the two approaches provided congruent results and independently revealed the following dominant patterns of introgression (Fig. 4; Supplementary data 2, Figs S1, S2). Markedly elevated values of private alleles and read mapping efficiencies were detected in the Hyrcanian population (Hyr) in relation to *R. dolichocarpus*. This was also the case for the three Caucasian samples (Cau) from Kakheti (easternmost Georgia with the common occurrence of this diploid), but not for the other Caucasian individuals from western regions. Those, on the other hand, exhibited a significantly increased affinity to *R. moschus*. The affinity to *R. ulmifolius* agg. was decreased in the Asian populations, but to a lesser extent also in the Balkan, Carpathian and Hercynian populations (Bal, SEC, WCa, Her), whereas all the other populations exhibited higher values but also with higher variability. Increased affinity to *R. canescens* was detected in the populations Bal and Ape (both northern and southern), but also with rather high variability.

**Fig. 4:**
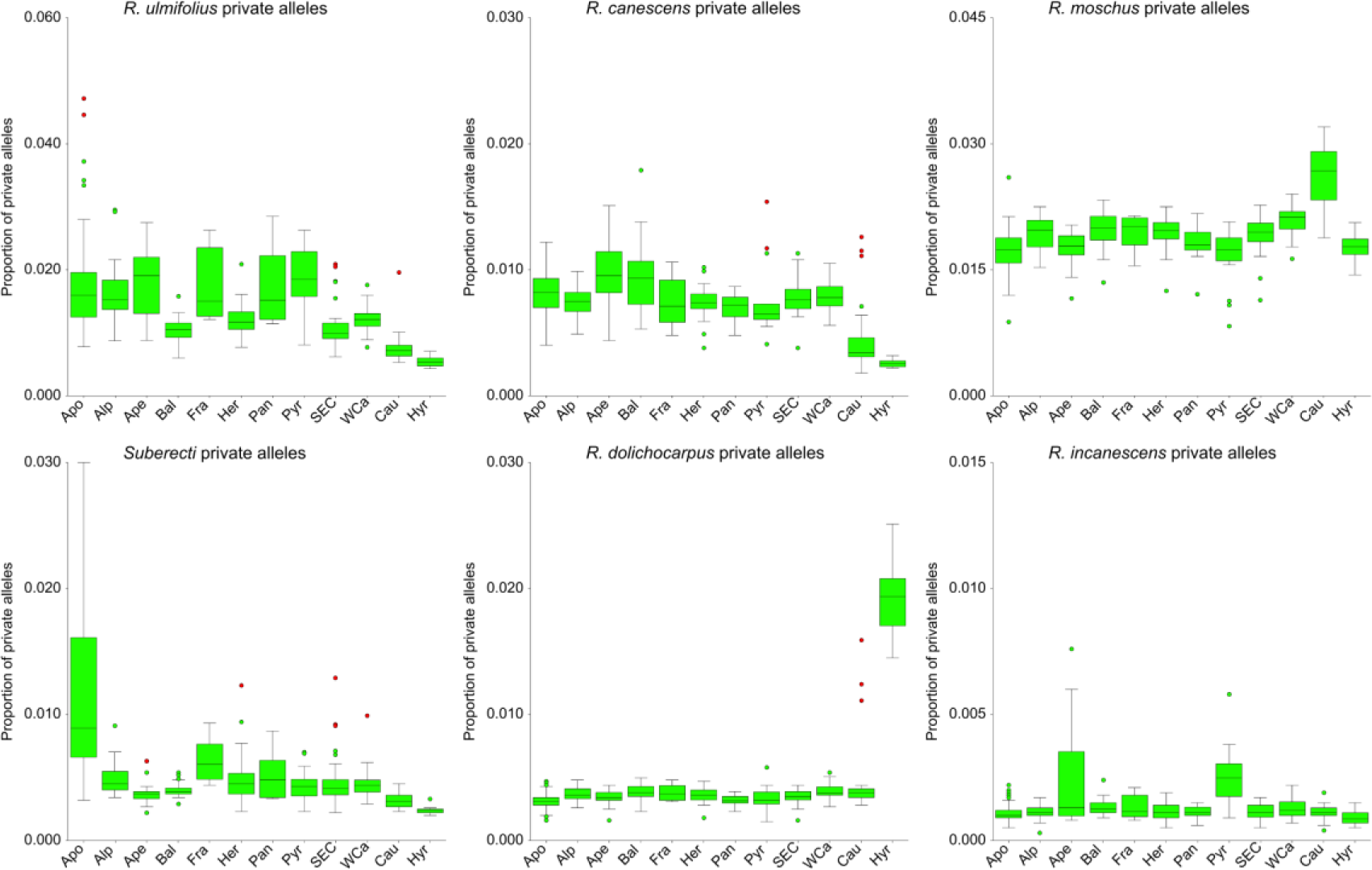
Proportion of private alleles of ancestral taxa detected in the *Glandulosi* populations: apomicts (Apo), Alps (Alp), Apennines (Ape), Balkans (Bal), France (Fra), Hercinia (Her), Pannonia (Pan), Pyrenees (Pyr), South-eastern Carpathians (SEC), Western Carpathians (WCa), Caucasus (Cau), Hyrcania (Hyr). For the full data set (including non-*Glandulosi* taxa, and *R. caesius* and *R. idaeus* private alleles), see Supplementary data 2, Fig. S2. Boxes indicate 25th, 50th and 75th percentile, whiskers show the largest/smallest observation that is less than or equal to the upper/lower edge of the box plus the 1.5× the box height; mild outliers (multiplier 1.5) shown as green dots, severe outliers (multiplier 3.0) in red.

Significantly increased affinity to the *Suberecti* was detected in apomicts and to a lesser extent in French sexuals (Fra) and three Pannonian (Pan) individuals, in the latter always in association with increased affinity to *R. ulmifolius* agg. Searching for genetic traces of *R. incanescens* was slightly hindered by the availability of only four individuals from two localities of this species for analysis. However, increased proportions of its private alleles were detected in almost all north-Apennine (Ape) individuals and also in most of the Pyrenean (Pyr) individuals. Increased affinity to *R. caesius* was only detected in three apomictic (Apo) individuals. Similarly, the affinity to *R. idaeus* was generally very low and with only a few outliers of unclear interpretation.

### 3.4. Population structure and coancestry matrices

STRUCTURE analysis resulted in the best-supported K=4 in both of the sample sets (*Basic set*, *European set*), but K=5 (the second best) provided a biologically more reasonable interpretation in the *Basic set*. All variants of the analysis distinguished clusters corresponding to the diploid taxa and the *Glandulosi* and *Suberecti* ancestors, the last of which was detected mainly in *Glandulosi* apomicts, in much lesser degree in sexuals (mainly population Her; Fig. 5). Small affinities to *R. ulmifolius* agg. were detected in populations Alp, Ape (mainly its southern part) and Pyr. Low levels of signal from *R. canescens* were revealed mainly in Ape and Bal, to a lesser extent also in Pyr and SEC. On the other hand, the signal from *R. ulmifolius* agg. and *R. canescens* was mostly negligible in apomicts (except for ca 7 individuals). A clear coancestry was detected between *R. dolichocarpus* and the Hyr population (plus three Cau individuals from eastern Georgia). *Rubus moschus* clustered with polyploids of *R.* ser. *Glandulosi*; only for K=7 it formed a separate cluster partly shared with the Cau and Hyr populations (Supplementary data 2, Fig. S3).

**Fig. 5:**
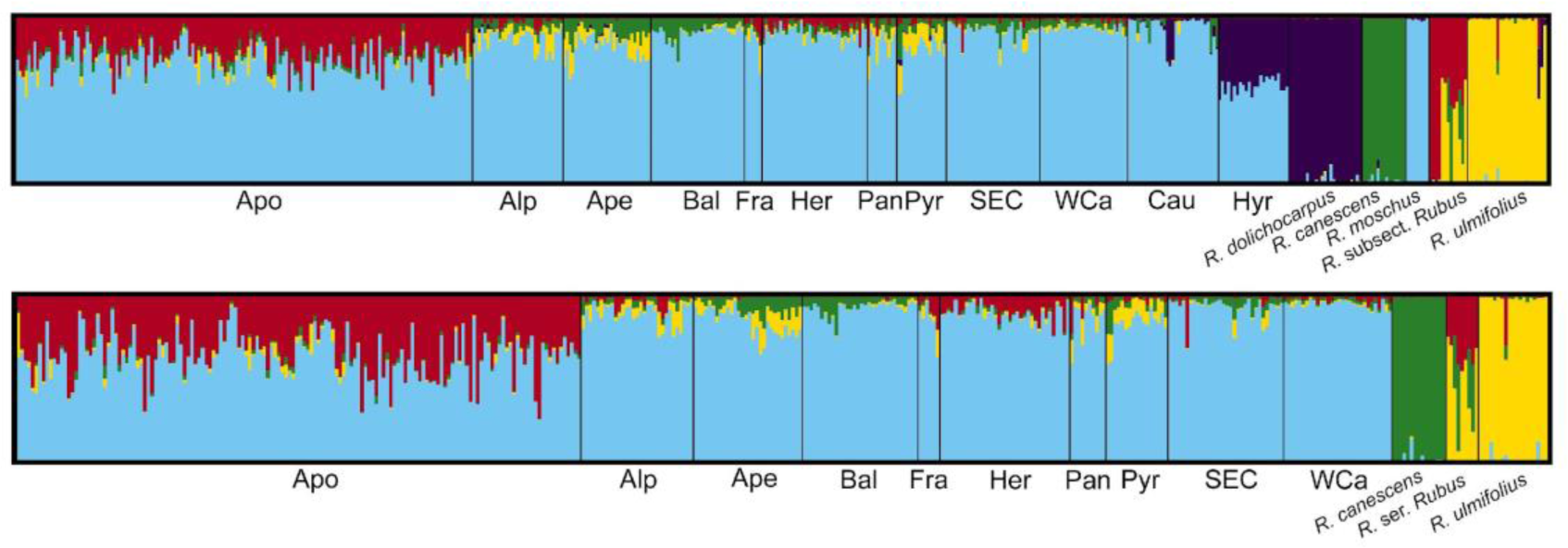
Population inference from STRUCTURE based on *Basic sample set* (above; 7,329 unlinked SNPs, K=5) and *European set* (below; 7,596 unlinked SNPs, K=4); each of the plots represents an average of ten runs (similarity score = 1.0). *Rubus* subsect. *Rubus* (above) covers both North American and European taxa (*R.* ser. *Rubus*) in this order. For the other K values, see Supplementary data 2, Figs S3 and S4.

The FINERADSTRUCTURE analyses indicated elevated coancestry mainly between Hyrcanian (Hyr; plus three Caucasian, Cau) sexuals and *R. dolichocarpus*, and between Cau sexuals and *R. moschus*. Three apomictic *Glandulosi* clearly showed affinity to *R. caesius,* slightly elevated coancestry was also detected between Pyr and Ape and *R. incanescens*, and between part of *Glandulosi* apomicts and *R.* subsect. *Rubus* (and other non-*Glandulosi* apomicts; Supplementary data 2: Fig. S5). Depending on parameters for data filtering, sample selection and FINERADSTRUCTURE settings, the tree-based population assignment (and thus the grouping of samples in heatmaps) was rather unstable, probably due to the reticulate nature of the studied system, but the resulting coancestry signal remained very similar across different analysis setups. Populations Cau and Hyr formed two separated groups from the European *Glandulosi* (yet with higher affinity to that group than to polyploid outgroups). They were, therefore, removed together with outgroups to reduce complexity.

In the resulting *European Glandulosi set*, most of the sexuals formed a separate group (Fig. 6), except for most of the Her (and in some replications also WCa; not shown) individuals, which exhibited increased affinity both to other *Glandulosi* sexuals and apomicts and their position was unstable. Within *Glandulosi* sexuals, a small, geographically diverse group of individuals exhibited a lower affinity to the other *Glandulosi* sexuals; this group also included three samples of *Glandulosi* apomicts and showed an increased affinity to *R. ulmifolius* agg. and non-*Glandulosi* apomicts (Supplementary data 2, Fig. S5), which most likely resulted from introgression from non-*Glandulosi* apomicts (marked in Fig. 6). The other sexuals were rather homogeneous and were constantly divided into two weakly differentiated clusters – the Balkan and Carpathian populations (Bal, WCa, SEC), and the western populations (Alp, northern Ape, Pyr, Fra). The southern part of the Ape population clustered constantly with the introgressed individuals probably due to their coancestry with *R. ulmifolius* agg., and exhibited higher affinity to Bal and SEC and the western populations (Alp, Pyr, Fra).

**Fig. 6:**
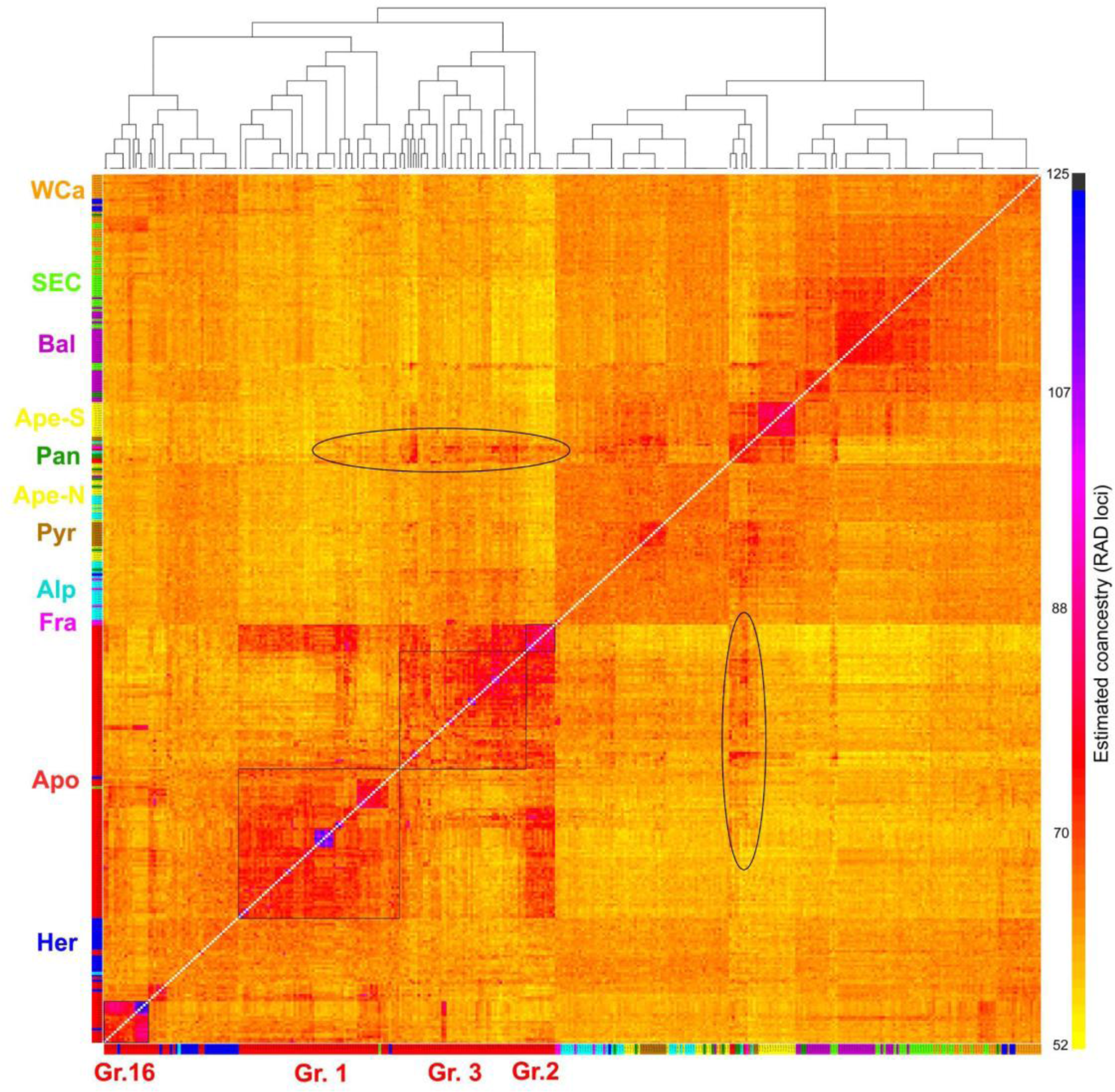
Coancestry matrix visualized as a heatmap based on the *European Glandulosi set* and 43,444 loci with 102,621 SNPs. Populations’ colours correspond to Fig. 2. The four groups of apomicts are marked as thin black squares, increased coancestry likely resulting from introgression from non-*Glandulosi* apomicts is marked in ovals.

*Glandulosi* apomicts, together with Her sexuals, tended to form four dominant, yet not always clearly delimited clusters. The most distinct one was the genotype 16 and its derivatives (Group 16), whose affinity to other apomicts was rather low. Group 1 apomicts shared its coancestry mainly with the Her sexuals and many individuals also with the WCa (and SEC and Bal) sexuals. Group 3 apomicts exhibited slightly elevated coancestry with the western sexuals Alp, Fra and Pyr. Apomicts of Group 2 exhibited markedly elevated coancestry with the Group 1 and 3 apomicts, but their affinity to *Glandulosi* sexuals (except for Her) was lower. Several individuals within Group 1 exhibited an elevated affinity to the Group 3, and two individuals from Group 3 exhibited marked coancestry with the genotype 16 in all analyses. Virtually the same results were obtained even after removing loci that contained private alleles of the ancestral species from analyses (not shown).

### 3.5. Present-day and paleodistribution modelling

The MAXENT model of the present-day predicted distribution of tetraploids of *R.* ser. *Glandulosi* with parameters LQHP with regularization parameter 1.0 was selected as the best in the model evaluation, based on the lowest AICc. Statistical validation suggested good model performance (AUC train = 0.838, AUC valid = 0.820±0.016, AUC diff = 0.021±0.016, OR10p = 0.127±0.035, mean±SD). The mean AUC for the ten replicated runs was 0.779±0.057. The highest mean relative contributions to the replicated models based on percent contribution and permutation importance, respectively, were provided by Bio5 (Max temperature of warmest month; 33.1%/25.9%), Bio6 (Min temperature of coldest month; 26.8%/18.7%), Bio12 (Annual precipitation; 21.8%/20.7%) and Bio15 (Precipitation seasonality; 12.3%/26.3%) variables. On the other hand, Bio2 (Mean diurnal range) and Bio3 (Isothermality) variables contributed little (always < 6%) to the models. Response and marginal curves of each bioclimatic variable are in Supplementary data 2: Fig. S6.

Mean MAXENT model of the predicted distribution under the current climate fitted quite well with the real distribution of *R.* ser. *Glandulosi*. Only westernmost Europe was slightly underestimated, probably due to the undersampling of this region, and the Baltic region was overpredicted compared to the real distribution of the tetraploids (Fig. 7a, b). The LGM models (Fig. 7e, f) predicted a suitable climate for the studied group almost exclusively in southern Europe (Iberian Peninsula, southern France, Apennine Peninsula, western and southern Balkans), western Asia Minor, along the southern and eastern Black Sea coast, and in Hyrcania. A comparison of two climatic scenarios of the LGM showed some regional differences in the predicted distribution mainly in Western Europe, western Asia Minor and the south-eastern Balkans. The Mid-Holocene models (Fig. 7c, d) predicted a suitable climate mainly in Central Europe north of the Alps, but a somewhat more patchy distribution of suitable climate was predicted also elsewhere in Europe and West Asia. Only small patches were predicted in the Apennine Peninsula, Crimea and Hyrcania.

**Fig. 7:**
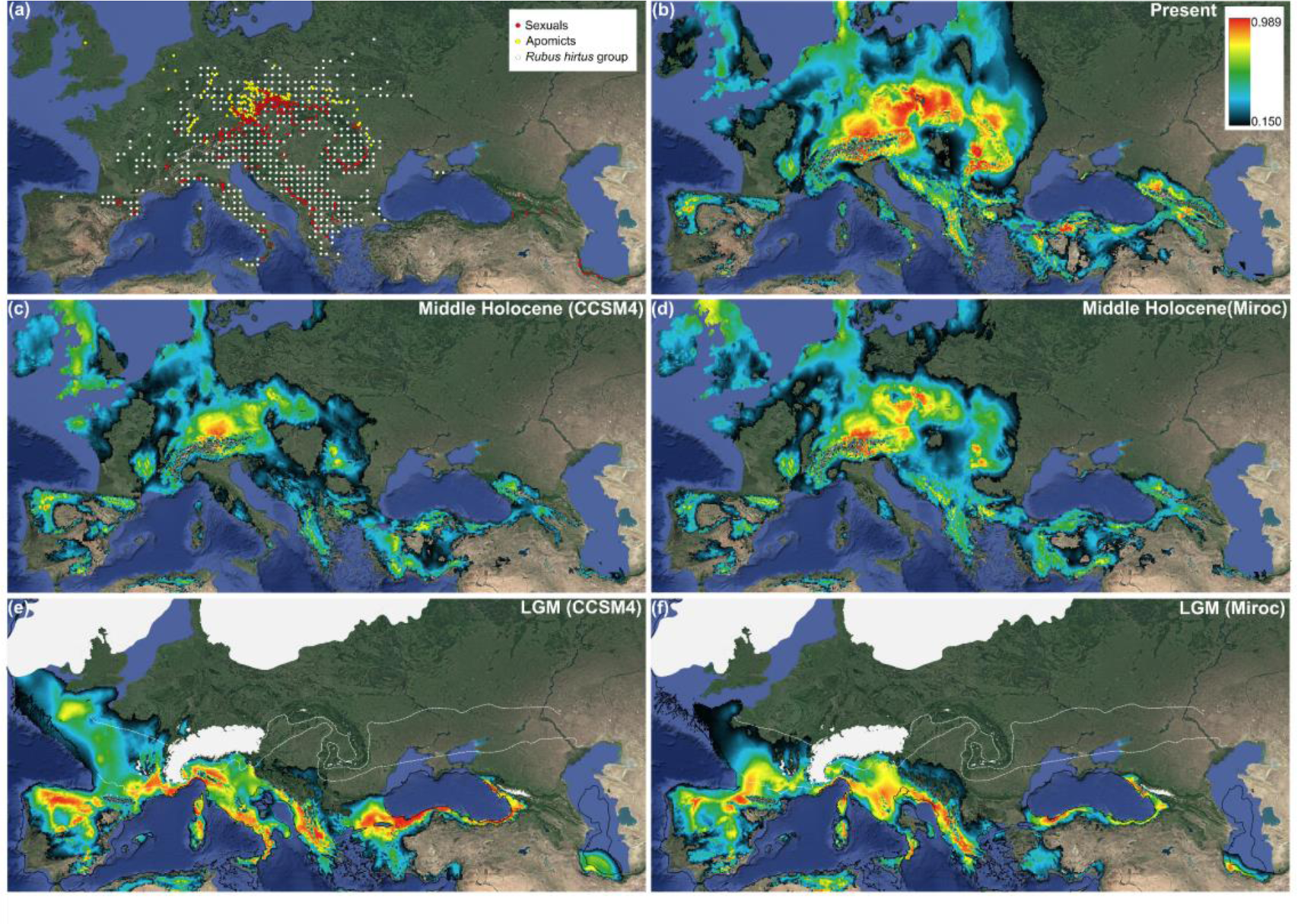
Distribution maps and climatic models. (a) Known distribution of *Rubus hirtus* group (Kurto et al. 2010) and localities of tetraploid apomicts (yellow points) and sexuals (red points) of *Rubus* ser. *Glandulosi* as published in Sochor et al. (under review). (b–f) Mean MAXENT models of predicted distributions (clog-log output, mean of 10 models each) for *R.* ser. *Glandulosi* (both apomictic and sexual) for the present, the mid-Holocene (6,000 years BP), and the Last Glacial Maximum (LGM; 21 ky BP). For past climates, projections on two climatic models are presented (CCSM4 – c, e; and Miroc – d, f), based on a distribution model computed with present-day climatic data. For the LGM, ice sheets and glaciers (white polygons), continuous and discontinuous permafrost (white dash lines) and paleocoastlines (black lines) are shown.

## 4. Discussion

### 4.1. Sexuals are not genetically impoverished along the contact zone with apomicts

In our previous work (Sochor et al., under review), we concluded that the extraordinary borderline between tetraploid sexuals and apomicts of *R.* ser. *Glandulosi* likely resulted from phylogeographic history and neutral processes because vast ecological niche overlap of sexuals and apomicts did not imply any role of (mal)adaptive factors in their geographical differentiation. However, the role of population genetic processes behind the observed spatial pattern, tentatively suggested already by Šarhanová et al. (2017), could not be ruled out. It is commonly presumed that marginal sexual populations tend to be small, fragmented, and impoverished in their genetic diversity (cf. Sagarin & Gaines, 2002; Eckert et al., 2008; Angert et al., 2020). This can lead to decreased adaptability, susceptibility to genetic drift, inbreeding or outbreeding depression (the latter usually due to secondary contact with related taxa), mutational meltdown, etc. (Keller and Waller, 2002; Willi et al., 2013). Additionally, continuous gene flow from central to marginal populations can prevent local adaptations (García-Ramos & Kirkpatrick, 1997). Due to their asexual reproduction, polyploidy and hybridity, apomicts can overcome or buffer such processes, and gain evolutionary advantage over the sexuals. Our RADseq did not reveal any differences among apomicts, sexual populations adjacent to the apomicts (mainly Her, WCa, Alp), and those further from their main range (Bal, Ape, Pan, SEC, Pyr) both in heterozygosity and inbreeding coefficient (Table 2). Furthermore, contrary to the expectation, the genetic diversity of individual genotypes (H_exp_) correlated negatively with the distance from the range of apomicts (Fig. 3). This likely implied a gene flow from *Glandulosi* apomicts to sexuals rather than genetic impoverishment of the latter, and reflected ecological optimum of sexuals (Fig. 7) and large sizes and high density of their populations in Central Europe (the authors’ personal observation). Therefore, our data do not support the view that population genetic processes typical for genetically impoverished populations can drive the geographic parthenogenesis in *R.* ser. *Glandulosi*.

### 4.2. Rubus ser. Glandulosi is introgressed by several taxa

*Rubus* ser. *Glandulosi* has always been regarded as one of the taxonomically most problematic *Rubus* taxa (Kurtto et al., 2010). This fact stems not only from varying levels of sexuality, recently characterized as geographic parthenogenesis (Sochor et al, under review), but also from an extreme morphological variation and plasticity (“amorphous” taxon; Holub, 1997). Although a relatively low diversity of plastid haplotypes and ITS ribotypes was detected in the group in Europe (Sochor et al., 2015), a distinct haplotype was detected in West Asia (Sochor and Trávníček, 2016; Kasalkheh et al., under review), which may imply a genetic contribution of an unknown (and probably extinct) ancestor in that region. Besides that, we detected at least some traces of each of the known Eurasian *Rubus* ancestors (see Table 1) in our *Glandulosi* RADseq data, except for *R. idaeus*. The most prominent shared ancestry was detected between the *Glandulosi* apomicts and *R.* subsect. *Rubus* (*Suberecti* ancestor; see below), and between *R. dolichocarpus* and *Glandulosi* sexuals across the range of *R. dolichocarpus*. *Rubus moschus* likely contributed to the tetraploid gene pool twice; firstly when it (or its predecessor) established the common tetraploid *Glandulosi* gene pool (light blue in Fig. 5; see also Sochor et al., 2015), and secondly in the later evolution in the Western Caucasus (Fig. 4; Supplementary data 2: Figs S1, S3). *Rubus incanescens* has hitherto not been suspected as an ancestor for polyploid brambles, probably due to its restricted distribution and rather sparse occurrence (Sochor et al., 2015). However, we have traced its alleles in *R.* ser. *Glandulosi* in the northern Apennines and the Pyrenees (Fig. 4), although in low quantities which did not enable a clear recognition of this ancestor in other analyses. It is, therefore, likely that *R. incanescens* has played some role in the evolution of the northwestern Mediterranean *Rubus* floras, yet its extent cannot be estimated at this moment. The two remaining Eurasian diploids, *R. canescens* and *R. ulmifolius* agg., exhibited regionally elevated genetic signatures in *R.* ser. *Glandulosi* (Figs 4 and 5). The last known *Rubus* ancestor, the tetraploid *R. caesius*, was traced only in three apomictic individuals and appears to be of low importance in the *Glandulosi* evolution, as we were otherwise unable to confirm its ancient (cf. Sochor et al., 2015) or contemporary involvement.

Although we detected a non-negligible gene flow from six diploid (including the presumably extinct one) species, its mechanism across ploidy levels remains unclear. The mechanism of triploid bridge (Ramsey and Schemske, 1998) appears to be rather insignificant as we have detected only one triploid hybrid of *R.* ser. *Glandulosi* and a diploid (*R. dolichocarpus*), and the hybrid was sterile, which was expected in a triploid hybrid of two sexuals. On the other hand, two putative hybrids with *R. ulmifolius* from Italy (with no other species detected at the localities) proved to be tetraploid (not shown) and indicated the possibility of hybridization via a reduced gamete of *Glandulosi* and an unreduced gamete of the diploid. Alternatively, the detected gene flow can proceed via non-*Glandulosi* polyploids, which frequently co-occur with *R.* ser. *Glandulosi*, produce viable hybrids (Šarhanová et al., 2012, 2017), and are themselves derived from the diploid ancestors (Sochor et al., 2015). This mechanism is presumed to be dominant, e.g., for the gene flow from *R. ulmifolius* via derived polyploids to Pannonian *Glandulosi* (Pan) for two reasons; (1.) *R. ulmifolius* agg. does not occur in that region (Kurtto et al., 2010), and (2.) the elevated frequency of *R. ulmifolius* agg. alleles in these individuals is associated with alleles of other ancestors, such as *R. canescens* and *Suberecti*. However, distinguishing between the two homoploid mechanisms is difficult in most other cases.

### 4.3. What is the origin of apomictic Glandulosi?

Our ddRADseq analysis showed that all *Glandulosi* apomictic genotypes share a part of their genomes—here called *Suberecti* subgenome—with each other but not with sexuals (or only to a minor extent). At the same time, this gene pool is shared with non-*Glandulosi* apomicts, particularly with *R.* subsect. *Rubus*, including the North-American taxa and *R.* ser. *Nessenses*. Both North-American taxa and *R.* ser. *Nessenses* are supposed to have undergone prolonged isolation from other European brambles, and are thought to share a common ancestor, here referred to as *Suberecti*, which is now extinct in Europe (see also Sochor et al., 2015). If this shared ancestry were the product of current hybridization of *Glandulosi* sexuals and non-*Glandulosi* apomicts, increased proportions of other ancestors’ alleles would also be expected in the *Glandulosi* apomicts. Apomicts from most taxonomic *Rubus* series, including ser. *Hystrix*, *Anisacanthi*, *Vestiti*, and *Pallidi*, indeed exhibit an increased genetic affinity to *R. ulmifolius* agg.—one of the main ancestral species (Supplementary data 2: Figs S1, S2). As mentioned above, the increased proportion of both *R. ulmifolius* agg. and *Suberecti* alleles was also detected in several sexuals or transitional individuals (sensu Sochor et al., under review), which may represent occasional hybrids or introgressants of the Holocene origin (marked in Fig. 6). On the contrary, most *Glandulosi* apomicts are remarkably similar to the sexuals in their low affinity to *R. ulmifolius* agg. (Figs 4 and 5). Therefore, our data imply that the *Suberecti* subgenome was not obtained by *Glandulosi* from the modern *Rubus* taxa, but probably from the now extinct *Suberecti* ancestor in the pre-glacial period. Subsequently, these *Glandulosi* apomicts formed a distinct phylogeographic lineage independent of the sexual lineage. The detected gene flow between the two lineages only results from their secondary contact. This scenario is also supported by the relative genotypic richness of apomicts, which is positively correlated with the distance from the contact zone with sexuals (Sochor et al., under review).

However, the authors’ personal observations suggest that the absolute number of apomictic genotypes is likely to be higher along the contact with sexual populations as a result of higher population densities. Consequently, it is expected that the sexual populations play a role in generating novel apomictic genotypes. Increased genetic affinity between apomicts and the adjacent sexual populations (Alp, Her, WCa) confirms this expectation (Fig. 6). However, coherence of *Glandulosi* apomicts in phylogenomic clustering implies that the gene flow from sexuals to apomicts is probably not the primary driving force in apomicts’ evolution. Interestingly, the STRUCTURE results indicate also the reverse gene flow to a lesser extent, i.e., from *Glandulosi* apomicts to sexuals, particularly in the Her population (Fig. 5).

Four groups can be tentatively distinguished in *Glandulosi* apomicts (Fig. 6). Group 3 has a very large distribution range (Supplementary data 2: Fig. S7) and includes also a few pentaploids (incl. *R. nigricans*). Group 2 has a narrow Central European range with just a single genotype in the West (determined as *R. serpens*/*angloserpens*). Group 1 has a Central to Eastern European distribution. The fourth group is formed around Genotype 16, which is possibly the most widespread tetraploid genotype in *R.* ser. *Glandulosi* (Sochor et al., under review). This genotype served apparently as an ancestor for many derived genotypes of smaller distribution, the other ancestors being Her/WCa sexuals or (in two detected cases) the Group 3 apomicts. It is unclear whether this grouping reflects their distinct origin (e.g., from isolated glacial refugia), or it is only artificial and mirrors different patterns of introgression. Considering the weak differentiation (Fig. 6; not detected in Fig. 5), the latter appears more likely. Group 2 shows strong coancestry to both Groups 1 and 3, but decreased coancestry to sexuals (except for Her), and may represent either “genetically pure” *Glandulosi* apomicts or hybrids of Groups 1 and 3. Group 1 is clearly related to Central European sexuals (Her and partly also WCa), whereas Group 3 has the highest affinity to western populations (Alp, Pyr).

### 4.4. Apomictic R. ser. Glandulosi may have migrated from two glacial refugia

We modelled a suitable climate for tetraploid *R.* ser. *Glandulosi* in two extreme time points in the recent geological past: the Last Glacial Maximum (LGM), and the mid-Holocene. During the first (cold and dry) extreme, *R.* ser. *Glandulosi* was probably restricted to southern refugia, similarly to other nemoral species, as the models correspond well to the modelled distribution, e.g., of *Fagus sylvatica* and *Carpinus betulus* (Svenning and Kageyama, 2008). Although the two used climatic models differ in predictions in several regions, *R.* ser. *Glandulosi* was likely able to survive the LGM in large continuous areas across southern Europe and in a more fragmented manner in West Asia (Fig. 7). Current distribution of *R.* ser. *Glandulosi* sexuals suggests their main LGM refugia in the Balkans and the Apennines/southern Alps. This is consistent with relatively low genetic differentiation (Figs 5, 6) and low numbers of private alleles in European populations compared to those in the Caucasus and Hyrcania (Table 2), which were (and still are) more isolated (Fig. 7). Highly suitable climate was also predicted in southern France along the southern margin of discontinuous permafrost (Fig. 7e, f). This region is the best candidate for the main glacial refugium of *R.* ser. *Glandulosi* apomicts due to the slightly increased coancestry of the apomictic Group 3 and the western sexuals (Alp, Pyr). These apomicts are widely distributed from France to Romania (Supplementary data 2: Fig. S7). Therefore, this group of likely western origin may be ancestral to Central and Eastern European apomicts. Its survival north of the Alps is unlikely (Müller et al., 2003; Binney et al., 2017; Janská et al., 2017) and unsupported by our climatic models. However, southern France may have provided suitable conditions during the LGM, as mesophilous tree species were recorded in the lower Rhone basin or the Pyrenean valleys (Delhom and Thiébault, 2005; Beaudouin et al., 2007) or even at somewhat higher latitudes in the Landes de Gascogne (Svenning et al., 2008; de Lafontaine et al., 2014). From these regions, apomicts may have spread along the Alps to Central Europe, which appears to have been the climatically most suitable region in warm and moist periods of the Holocene. In contrast, areas with a suitable climate became more fragmented and restricted to high mountains in southern Europe (Fig. 7c, d), similar to the current state. This scenario of apomicts’ spread, however, does not appear to be likely for Genotype 16, which shows a very low coancestry with the other apomicts and western sexuals, but exhibits a higher affinity to eastern sexuals—particularly several WCa and SEC individuals. Its extensive distribution area, covering the whole of Central Europe from Czechia to Romania, and its apparent role in formation of other apomictic genotypes suggest it is an old genotype of probably different phylogeographic history. Our models do not support its large-scale survival during the LGM in the Carpathians. However, the southern foreland of the Western Carpathians demonstrably hosted a woodland biota in protected and relatively moist valleys (Ložek, 2006; Jankovská and Pokorný, 2008). The same situation probably existed in the Eastern Carpathians with a possible regional survival of deciduous trees (Magyari et al., 2014; see also Molnár et al., 2023). The survival of one generalistic apomictic genotype is, therefore, plausible. Such a scenario would imply that *R.* ser. *Glandulosi* apomicts were widespread in Central Europe also in the previous interglacial period and later contracted into two (or more) glacial refugia. Because the area of possible survival of Genotype 16 is now inhabited by sexuals (Sochor et al., under review), this scenario would also necessarily imply that the contact zone is not stable and apomicts are systematically edged out by sexuals, e.g., by the mechanism of clonal turnover (Janko, 2014). Alternatively, Genotype 16 may have been formed *de novo* in the Holocene from Carpathian sexuals completely independently of the other *Glandulosi* apomicts. This is, however, unlikely without a contribution of non-*Glandulosi* apomicts, whose genetic footprints were not detected in our RADseq data (Supplementary data 2: Fig. S5), and neither are they suspected from morphology. At the same time, the *Suberecti* subgenome does not deviate between Genotype 16 and other *Glandulosi* apomicts (probabilities of assignment of Genotype 16 to the *Glandulosi*: *Suberecti* cluster being constantly around 86:14 with no other cluster assigned; Fig. 5).

## 5. Conclusions

*Rubus* ser. *Glandulosi* is a widely distributed taxon with a complex evolutionary history, which resulted in a specific pattern of geographic parthenogenesis. The whole group is introgressed from six species (or seven considering the unknown Caucasian ancestor inferred from plastid haplotypes, see paragraph 4.2), but the gene pool of a single, extinct species— *Suberecti*—is characteristic for the apomicts and appears to be determinative for their reproductive mode. A fast postglacial recolonization of Central and Western Europe by apomicts from the main western and possibly another, small eastern refugium, accompanied by migration of sexuals from the Balkans and the Apennines/southern Alps probably established the contact zone between the reproductive modes. How stable this line of contact is and which factors play a role in its shifts is unclear. As the available data do not suggest any differences in competitive abilities between sexuals and apomicts (Konečná et al., unpublished data), we hypothesize that sexuals should gradually extend their distribution into the range of apomicts due to the neutral process of clonal turnover, while the speed of this shift is determined by genotypic diversity and population density of apomicts (Sochor et al., under review). In such a case, fast recolonization of Central Europe from the west may have been crucial for establishing a relatively stable position of the apomicts that guarantees resistance against the expansion of sexuals.

## Author contributions

Design of the research: MS, with a contribution of PŠ. Performance of the research – sampling: MS, BT with a contribution of MD, MH, MK; labwork: MS, PŠ; data analysis: MS with a contribution of MK; modelling: MD. Interpretation: MS. Writing the manuscript: MS with a contribution of MD. All authors contributed to and approved the final version of the manuscript.

## Acknowledgements

We thank particularly the many sample contributors, namely Gergely Király, Martin Lepší, Petr Lepší, Michael Hohla, Abraham van de Beek, Friedrich Sander, Pavol Eliáš jun., Věra Forejtová, Konrad Pagitz, Adriano Soldano, Henri Michaud, Piotr Kosiński, Martin Dančák, and others. We also thank Razieh Kasalkheh and Saeed Afsharzadeh for their valuable help with sampling in Hyrcania. Jakub Slatinský is acknowledged for writing the script for private alleles’ tracking.

The work was financed by the Grant Agency of the Czech Republic (www.gacr.cz) as a project no. 21-01233S. PŠ was supported by Operational Programme Research, Development and Education – Project “Postdoc2MUNI” (No. CZ.02.2.69/0.0/0.0/18_053/0016952).

Computational resources were provided by the e-INFRA CZ project (ID:90254), supported by the Ministry of Education, Youth and Sports of the Czech Republic.

