## Supplementary data 2 for "Plant kleptomaniacs: geographic genetic patterns in the amphi-apomictic *Rubus* ser. *Glandulosi* (Rosaceae) reveal complex reticulate evolution of Eurasian brambles"

### Supplementary data 2; Fig. S1

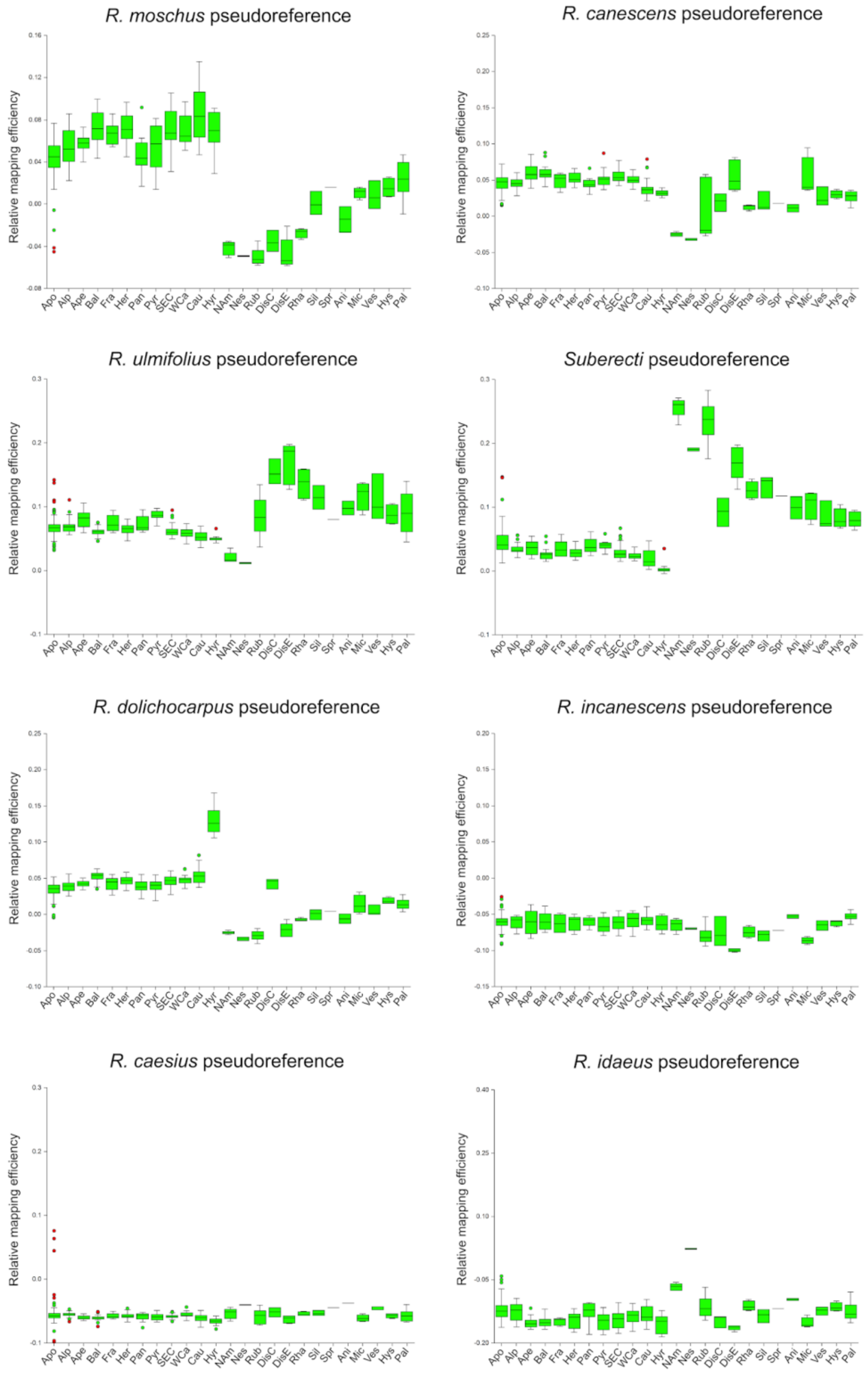


**Fig. S1:** Boxplots of relative mapping efficiencies of reads to psedoreferences of ancestral taxa. *Rubus* ser. *Glandulosi* populations: apomicts (Apo), Alps (Alp), Apennines (Ape), Balkans (Bal), France (Fra), Hercinia (Her), Pannonia (Pan), Pyrenees (Pyr), South-eastern Carpathians (SEC), Western Carpathians (WCa), Caucasus (Cau), Hyrcania (Hyr); outgroups: North American taxa (NAm), *R.* ser. *Nessenses* (Nes), *R.* ser. *Rubus* (Rub), *R.* ser. *Discolores* from the Caucasus and Europe (DisC, DisE, respectively), *R.* ser. *Rhamnifolii* (Rha), *R.* ser. *Silvatici* (Sil), *R.* ser. *Sprengeliani* (Spr), *R.* ser. *Anisacanthi* (Ani), *R.* ser. *Micantes* (Mic), *R.* ser. *Vestiti* (Ves), *R.* ser. *Hystrix* (Hys), *R.* ser. *Pallidi* (Pal).

**Fig. S2**

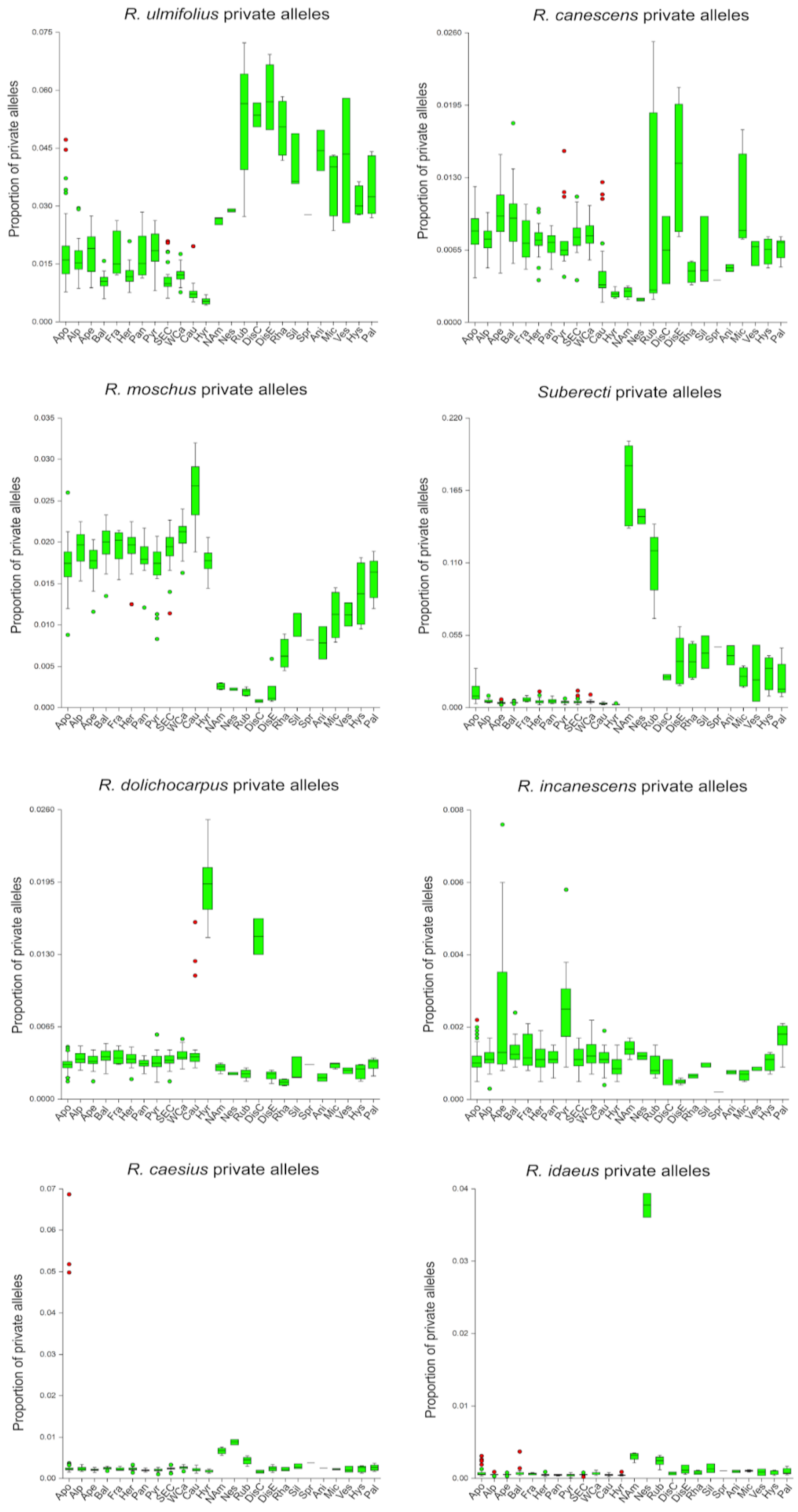


**Fig. S2:** Proportion of private alleles of ancestral taxa detected in *Rubus* ser. *Glandulosi* populations – apomicts (Apo), Alps (Alp), Apennines (Ape), Balkans (Bal), France (Fra), Hercinia (Her), Pannonia (Pan), Pyrenees (Pyr), South-eastern Carpathians (SEC), Western Carpathians (WCa), Caucasus (Cau), Hyrcania (Hyr); and outgroups – North American taxa (NAm), *R.* ser. *Nessenses* (Nes), *R.* ser. *Rubus* (Rub), *R.* ser. *Discolores* from the Caucasus and Europe (DisC, DisE, respectively), *R.* ser. *Rhamnifolii* (Rha), *R.* ser. *Silvatici* (Sil), *R.* ser. *Sprengeliani* (Spr), *R.* ser. *Anisacanthi* (Ani), *R.* ser. *Micantes* (Mic), *R.* ser. *Vestiti* (Ves), *R.* ser. *Hystrix* (Hys), *R.* ser. *Pallidi* (Pal).

**Fig. S3**

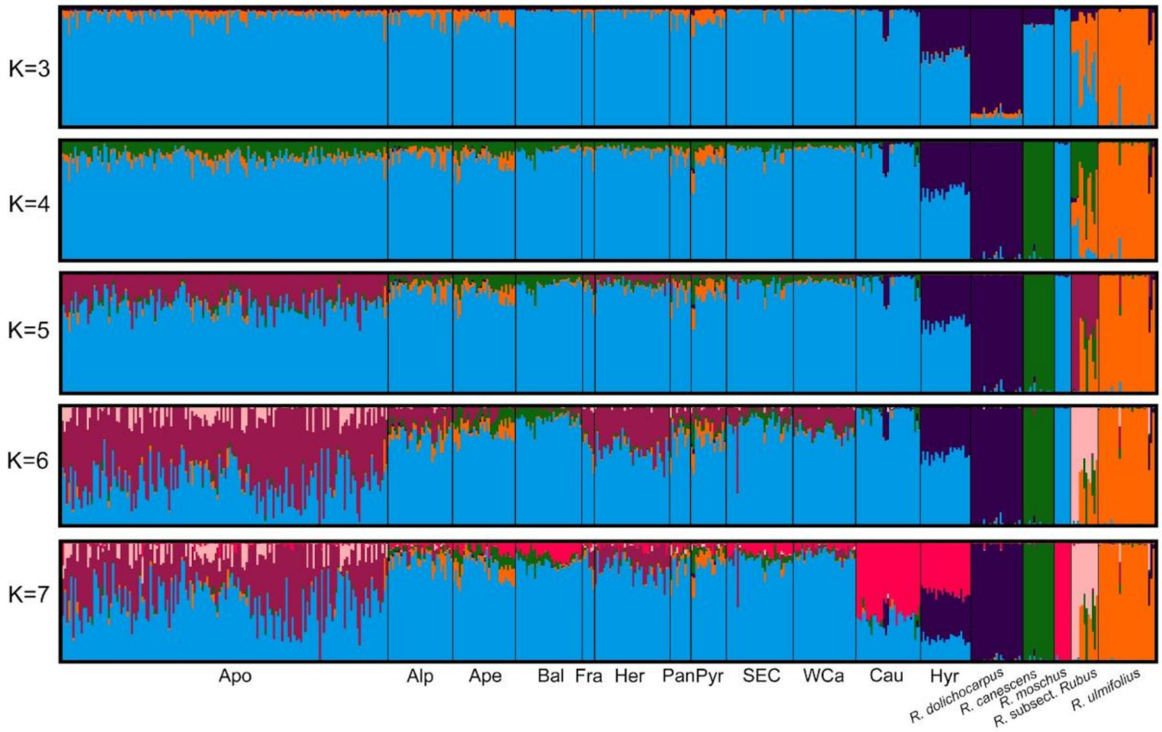


**Fig. S3:** Population inference from STRUCTURE based on *Basic sample set* and 7329 unlinked SNPs; K=3–7, averages of ten runs per K (similarity score = 1.0 for each K). *Rubus* subsect. *Rubus* (*Suberecti*) includes both North American and European taxa in this order.

**Fig. S4**

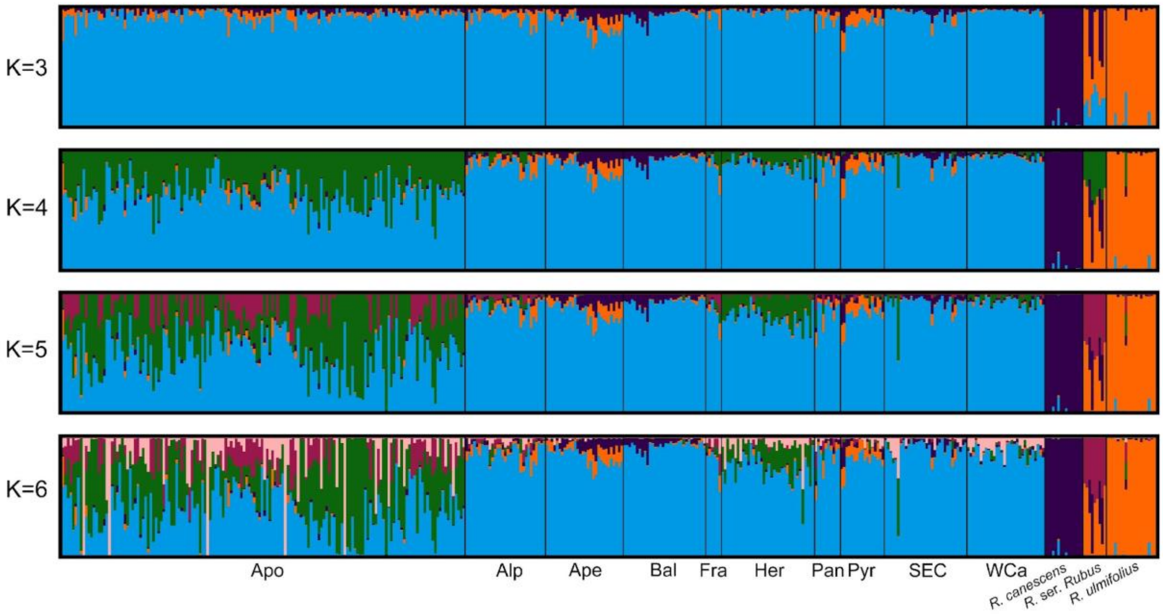


**Fig. S4:** Population inference from STRUCTURE based on *European dataset* and 7596 unlinked SNPs; K=3–6, averages of ten runs for K=3 to 5 (similarity score = 1.0) and six runs (major cluster) for K=6 (similarity score = 0.999).

**Fig. S5**

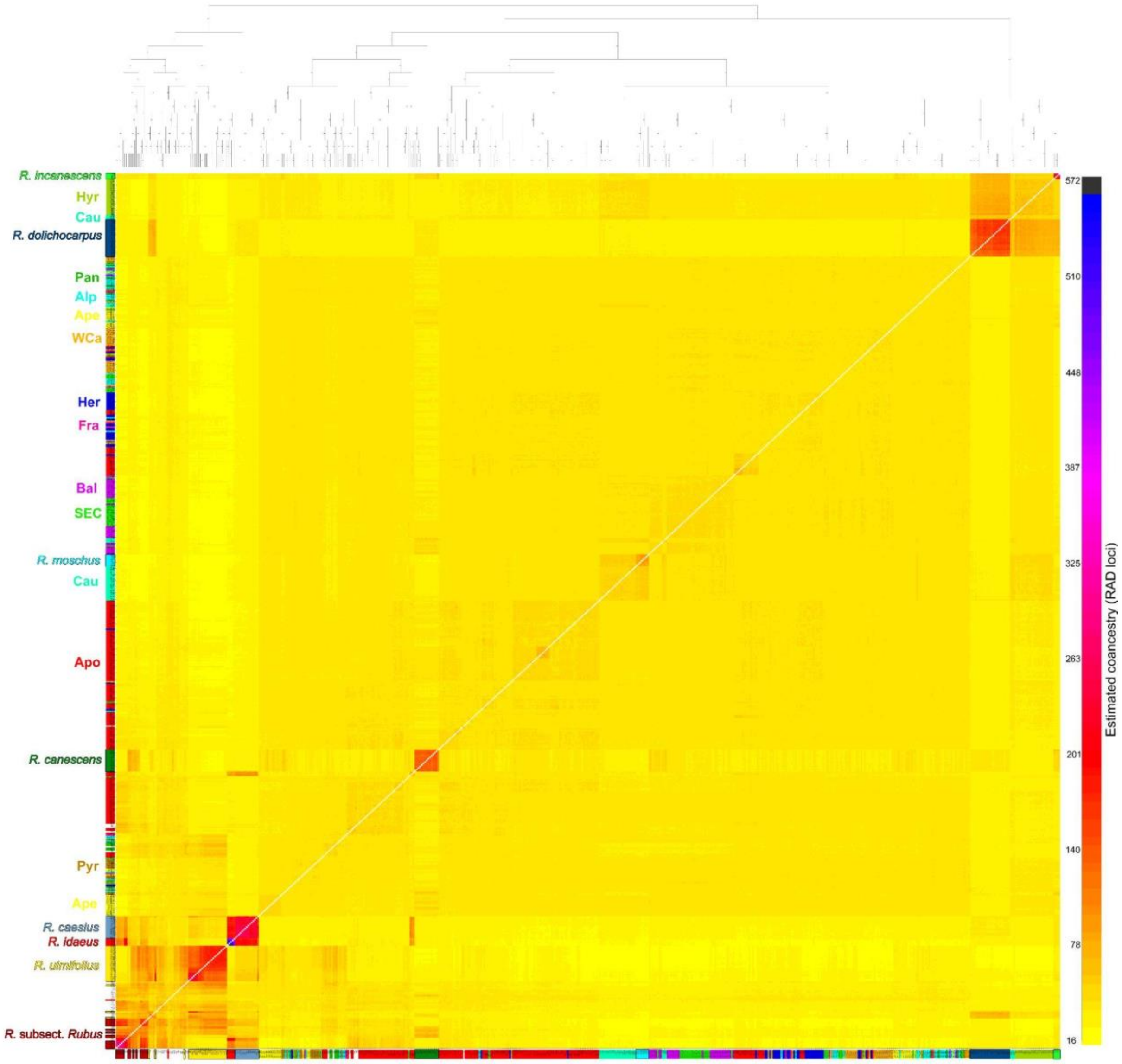


**Fig. S5:** Coancestry matrix visualized as a heatmap based on a complete sample set with 30,328 loci (98,362 variant sites). Populations' colours correspond to Fig. 1, derived polyploid outgroups other than *R*. subsect. *Rubus* (*Suberecti*) shown with white background.

**Fig. S6**

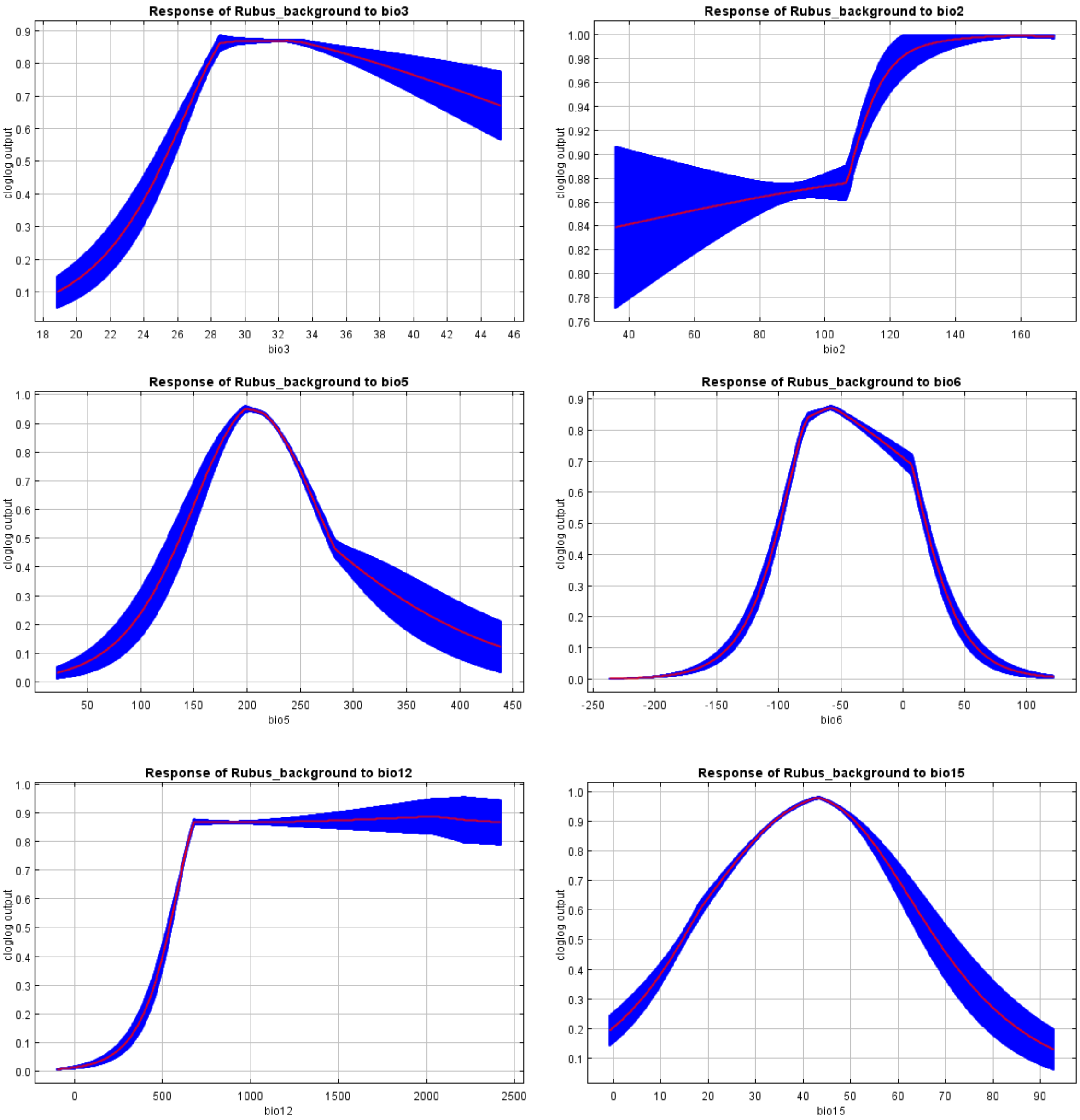


**Fig. 6A:** Response curves show how each environmental variable affects the Maxent prediction. The curves show how the predicted probability of presence changes as each environmental variable is varied, keeping all other environmental variables at their average sample value (mean+-1 SD from 10 runs).

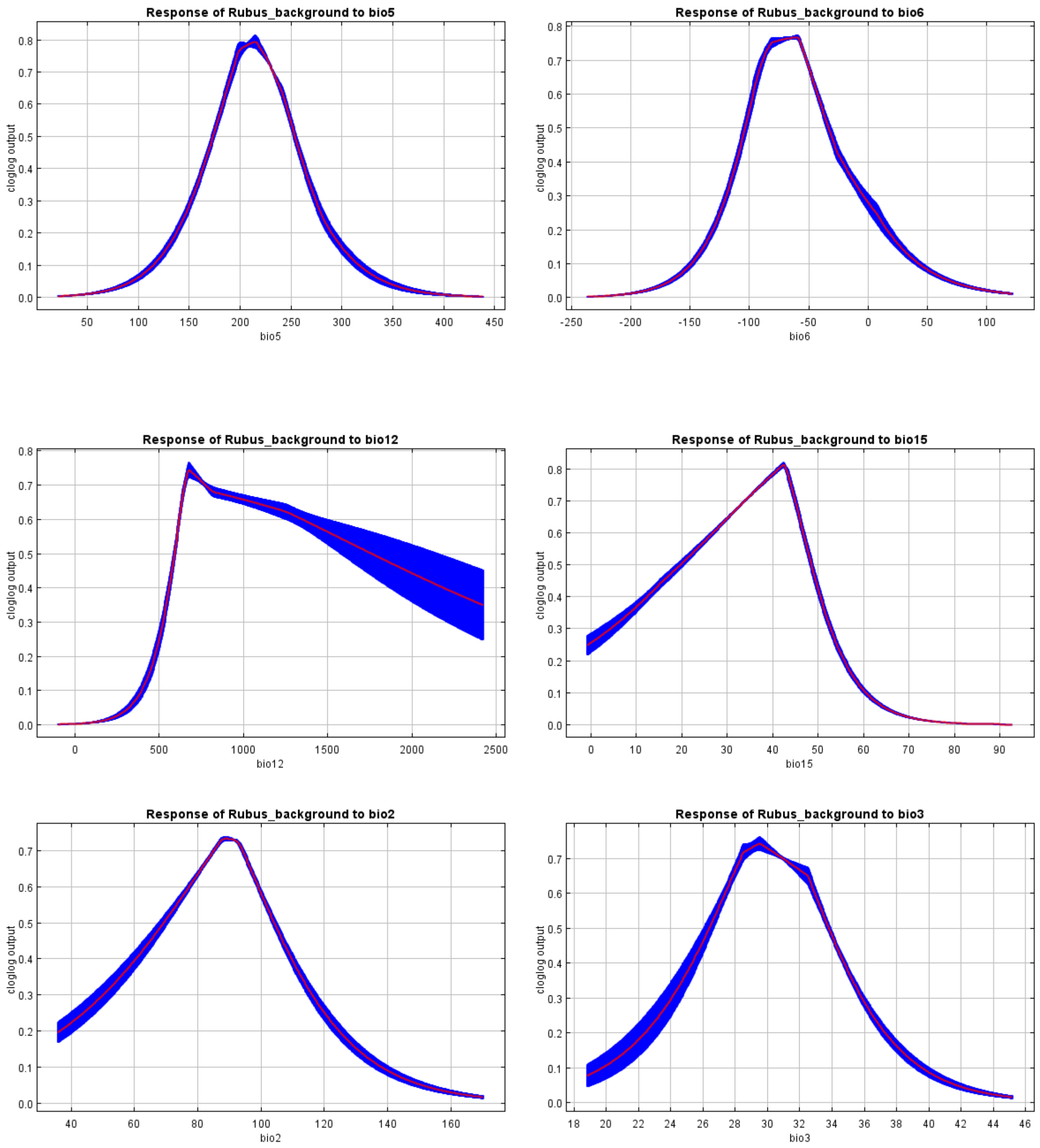


**Fig. 6B:** Marginal response curves; each of the curves represents a different model, namely, a Maxent model created using only the corresponding variable (mean+-1 SD from 10 runs).

**Fig. S7**

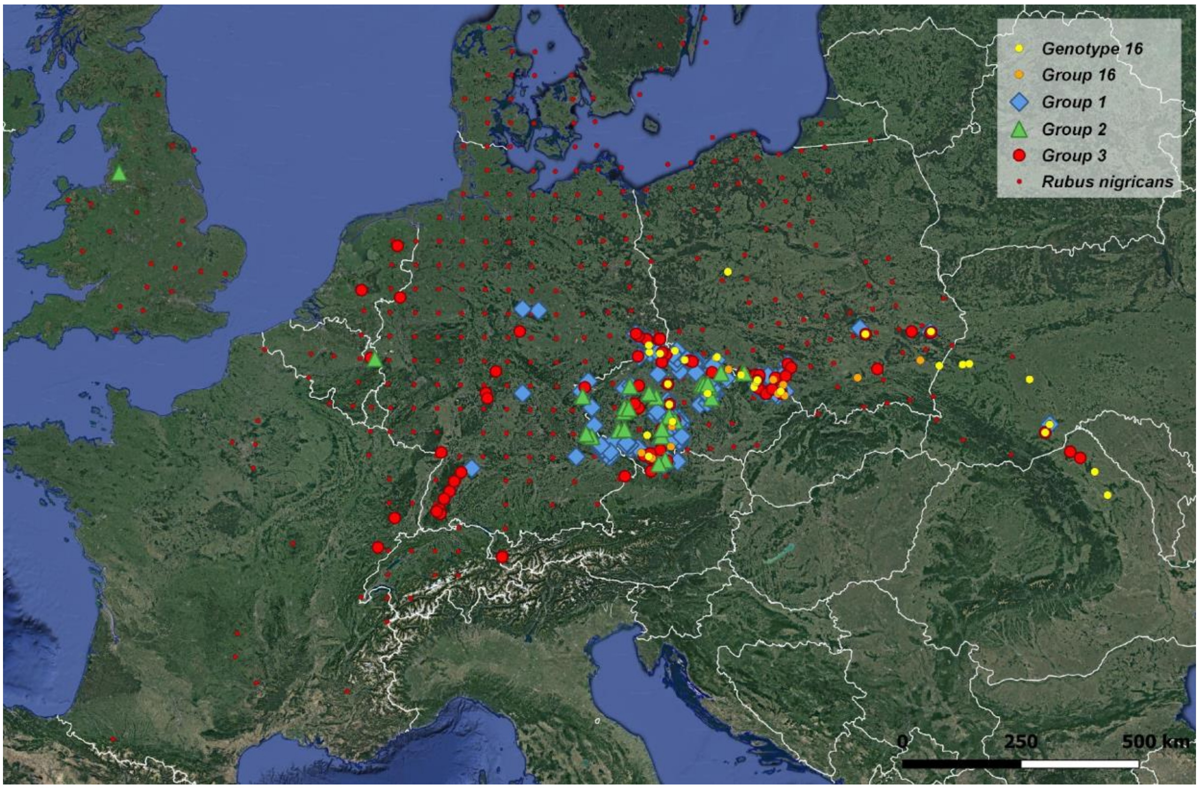


**Fig. S7:** Distribution of apomictic groups as defined by FINERADSTRUCTURE analysis. All samples used in Sochor et al. (under review) are included according to their genotypic identification; distribution of *Rubus nigricans* belonging to Group 3 is taken from Kurtto et al. (2010; under the name *R. pedemontanus*); genotype 16 is distinguished here from the rest of Group 16 by a different symbol.
